# Single-cell transcriptomic analysis identifies neocortical developmental differences between human and mouse

**DOI:** 10.1101/2020.04.23.056390

**Authors:** Ziheng Zhou, Shuguang Wang, Dengwei Zhang, Xiaosen Jiang, Jie Li, Ying Gu, Hai-Xi Sun

**Author notes:** Corresponding authors (H.-X.S.), (Y.G.).

## Abstract

**Background:** The specification and differentiation of neocortical projection neurons is a complex process under precise molecular regulation; however, little is known about the similarities and differences in cerebral cortex development between human and mouse at single-cell resolution.

**Results:** Here, using single-cell RNA-seq (scRNA-seq) data we explore the divergence and conservation of human and mouse cerebral cortex development using 18,446 and 7,610 neocortical cells. Systematic cross-species comparison reveals that the overall transcriptome profile in human cerebral cortex is similar to that in mouse such as cell types and their markers genes. By single-cell trajectories analysis we find human and mouse excitatory neurons have different developmental trajectories of neocortical projection neurons, ligand-receptor interactions and gene expression patterns. Further analysis reveals a refinement of neuron differentiation that occurred in human but not in mouse, suggesting that excitatory neurons in human undergo refined transcriptional states in later development stage. By contrast, for glial cells and inhibitory neurons we detected conserved developmental trajectories in human and mouse.

**Conclusions:** Taken together, our study integrates scRNA-seq data of cerebral cortex development in human and mouse, and uncovers distinct developing models in neocortical projection neurons. The earlier activation of cognition -related genes in human may explain the differences in behavior, learning or memory abilities between the two species.

## Background

The mammalian cerebral cortex develops via a complex process of cell proliferation, differentiation, and migration events. The molecular features of the human brain at gestational weeks 7-23 are similar to those of the mouse at postnatal days 14.5 to birth (P0) and reveal gene expression differences between the two species [1, 2]. Interneurons and projection neurons, which are the two major classes of neurons, populate the neocortex. Interneurons show largely GABAergic, connecting locally within the neocortex, and are generated by progenitors in the ventral telencephalic (inhibitory cortical interneuron producing) radial glia, before migrating from sub pallial ganglionic eminences into the cortex [3-6]. By contrast, projection neurons are excitatory, which are generated by progenitors in the dorsal telencephalic (excitatory cortical neuron producing) radial glia [7-9].

Despite immense efforts over the past decades to study the cell types, function, differentiation for the mammalian cerebral cortex, especially human and mouse, large differences and conservation are observed in human versus animal models [10-12]. Elucidating the conserved, divergent and cellular architecture of developing the cerebral cortex, especially projection neurons developments, may highlight to understanding susceptibility to neurological diseases and the occurrence of cognition and memory abilities in the early stages.

Recently, with the advance of next-generation sequencing (NGS) techniques, single-cell RNA sequencing (scRNA-seq) technologies has been paved the way for study cell phenotype and cell behavior at single-cell resolution [13, 14]. More significantly, scRNA-seq can overcome the heterogeneity of cross-species analysis, bypassing the need for pure cell sorting strategies [15, 16]. Moreover, although many studies have been reported the single species cerebral cortex using scRNA-seq, there is no systematic research to integrate the human and mouse scRNA-seq data, and compare the conservation and divergence of developing neocortex.

Here we comprehensively characterize the conservation and divergence of the cerebral cortex program between human and mouse [9, 17, 18]. We present a cross-species cell atlas and in which no obvious differences in cell type composition was observed. However, we identified different developing trajectories of projection neurons that in human the subventricular zone (SVZ) migration to deeper neocortical layers and more superficial layers are two independent processes. Additionally, the deeper and upper layers are more precisely differentiated in human than those in mouse, and lack of neuron migration signal in human. By comparing excitatory neurons gene expression across species, we also observed significant expression differences of function genes for cell migration, cognition, learning and memory in humans compared with mice. We next evaluated whether further refinement of neuronal subtypes occurs as neurons mature, indicating that human neuronal subtypes resolve from coarse laminar distinctions to refined transcriptional states in the later development, whereas mouse may not show precise subtype differentiation. In summary, we show the importance of cross-evolutionary comparison to better characterize the differences of human and mouse projection neurons development, and the relevance of these to cognition pathway. In conclusion, our study offers essential clues for researching neurodevelopment, especially for the projection neurons development, and the differences of occurrence of cognition ability in human and mouse.

## RESULTS

### Integration of single-cell transcriptomics of the human and mouse cerebral cortex

In order to compare cellular and molecular processes of the late corticogenesis between human and mouse, we collected published scRNA-seq data sets of human and mouse cerebral cortex at different development stages [9, 17, 18]. We reanalyzed the transcriptome profiles of 18,446 mouse neocortical cells at two key time points for corticogenesis development including 10,836 cells at embryonic day 14.5 (E14.5) from six biological replicates (Fig. 1A), and 7,610 cells at birth (P0) from three biological replicates (Fig. 1A). Because 7-to 23-post-conception-week (pcw) human brain cells are analogous to E14.5 to P0 mouse brain cells (Fig. 1A) [1, 2], we incorporated the transcriptome profiles of 6,655 human brain cells including 100 cells between 5.85 and 7 pcw, and 800 cells between 23-37 pcw (Fig. 1A). To ensure that cells from different species and data sets can be properly integrated, we performed unbiased clustering using Uniform Manifold Approximation and Projection (UMAP) [19]. The major cell lineages in cerebral cortex such as excitatory neurons, inhibitory neurons, neuronal progenitor cells and glial cells were able to be identified using their lineage markers [20, 21] (Fig. 1B, C). Moreover, there was no obvious bias because the majority of the cell types could be identified in all three data sets (Fig. S1C).

**Figure 1.**
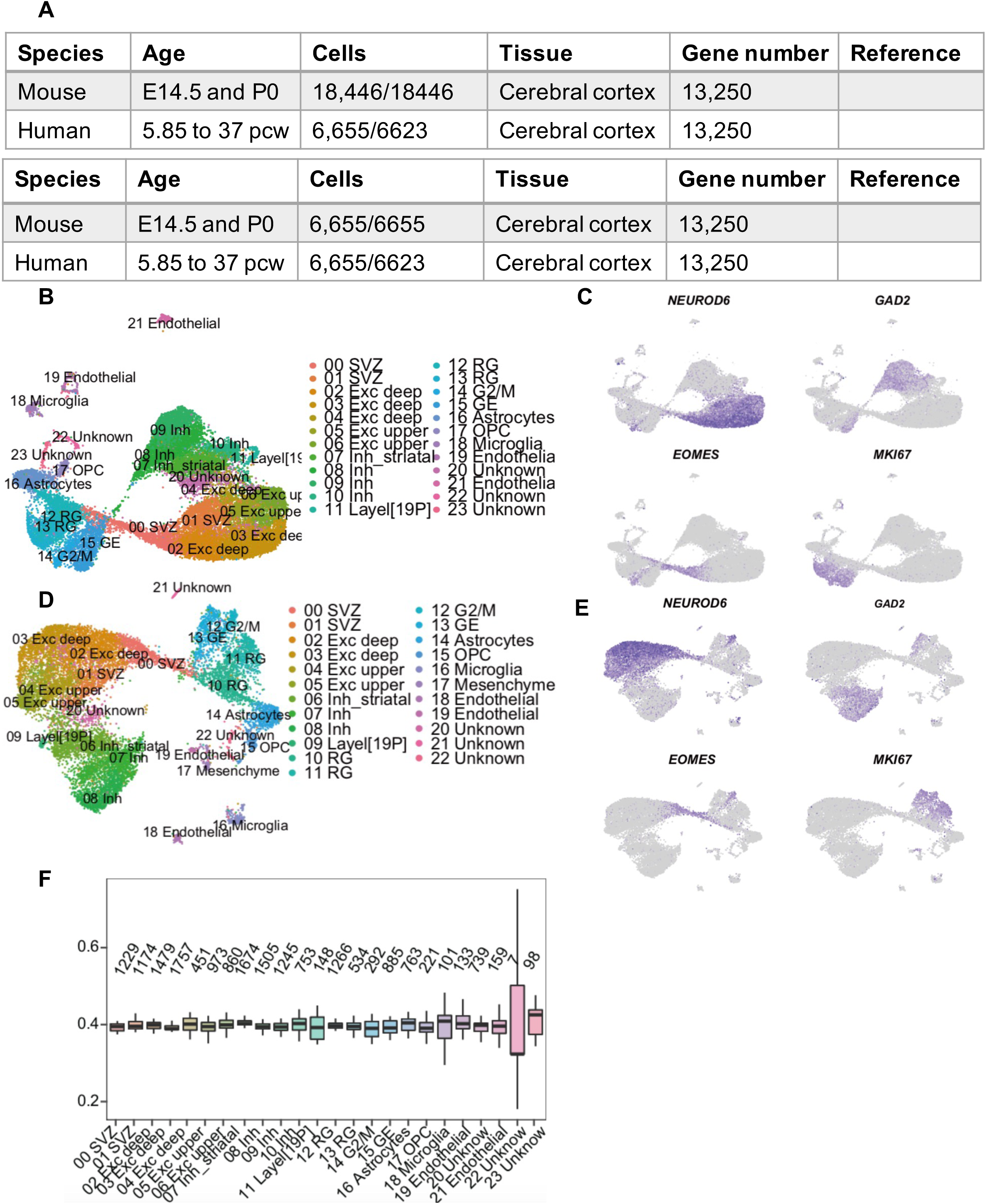

Because the number of mouse cells (18,446) was about 3 times higher than that of human cells (6,655) which may cause bias, we randomly extracted 6,655 mouse cells in further analysis. Finally, 13,278 cells (6,655 mouse cells and 6,623 human cells) were used (Fig. 1A) in further analysis which covers all major cell types (Fig. 1D, E).

### Identification and characterization of cortical cell types

We first compared the cell composition and found that mouse had more interneuron than human (Fig. 2A). We next compared the cell number in each cluster and found that cluster 06 Inh_striatal and cluster 09 Layer [19P] had more cells in mouse (20%-30%) than in human (70%-80%). In contrast, cluster 16 Microglia and 17 Mesenchyme had more human cells (80%-90%) (Fig. 2B).

**Figure 2.**
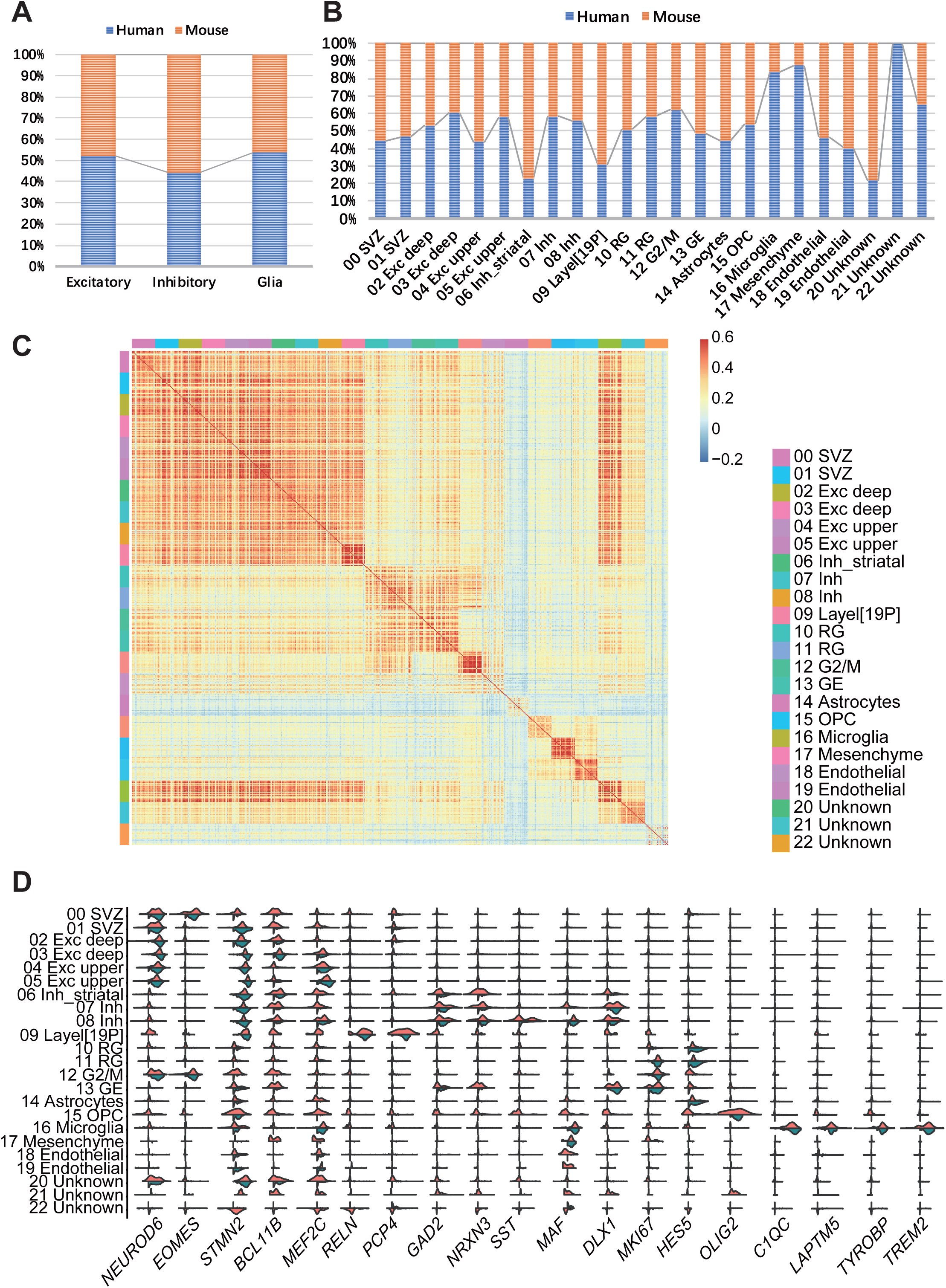

We next examined the correlation among clusters by cross-species clustering (Fig. 2C). As there were still considerable species differences between humans and mice, we used the integrated matrix. Remarkably, excitatory neuron exhibited a profile highly similar to interneuron, and Glia showed itself a high degree of correlation (Fig. 2C). Interestingly, Cluster 16 Microglia were distinct from excitatory neuron, interneuron and other Glia cells (Fig. 2C). We also calculated the correlation using human and mouse cells respectively and found that human and mouse had similar results with cross-species (Fig. S2A, B).

We checked the top 10 highly expressed genes of each cluster and found most of them were consistent among human and mouse (Fig. 2D, S2C, D). For example, two SVZ clusters (Clusters 00 SVZ and 01 SVZ) were featured by predominant expression of *EOMES* (*TBR2*) and *ZBTB20* [22-24]. Cells in Cluster 01 SVZ also expressed markers of both excitatory (*NEUROD6*) and inhibitory (*CALB2*) neurons [25-27] which exhibited the expression patterns of rostral migratory steam [18]. Cells in deep-layer populations (Clusters 02 Exc deep and 03 Exc deep) exhibited high expression levels of *BCL11B* [28]. Cells in Clusters 04 Exc upper and 05 Exc upper expressed upper layer marker *MEF2C* [8, 29]. We identified *NEUROD6* and *NEUROD1* kept consistently high expression in excitatory inhibitory, and *EOMES* expressed in subventricular zone (SVZ), and *STMN2* expressed strongly in deep and upper layer and increased the levels of expression with neuron maturation, which conversed the transformation known genes expression trajectories in neurogenesis among human and mouse (from *EOMES* to *STMN2*) [9, 30] (Fig. 2D). Three interneuron clusters (Clusters 06 Inh_striatal, 07 Inh and 08 Inh) were characterized by expression of *GAD2* and *NRXN3*. Specifically, in Cluster 08 Inh we found high expression of *LHX6, SST* and *MAF* indicating 08 Inh is somatostatin (SST) expressing interneurons but not parvalbumin (PV). *LHX6* is a transcription factor associated with PV and SST interneuron [31, 32], whereas *SST* and *MAF* regulate the potential of interneurons to interneurons to acquire SST+ interneuron identity [33]. Interneuron migrate tangentially from the ganglionic eminences, and ganglionic eminences (Cluster 13 GE) expressed high levels of inhibitory marker *DLX1* and proliferation markers *TOP2A* and *MKI67* [34]. RG marker *HES5* was in two RG clusters (Clusters 10 RG and 11 RG), while Cluster 11 RG expressed high levels of *MKI67* and *TOP2A* genes. We also found *GAP43*, a key regulator in early brain development, was constitutively expressed in all these cell types in both human and mouse [35].

We identified Clusters 06 Inh_striatal and 09 Layer[19P] as inhibitory neurons and they had similar gene expression patterns and cell numbers in human and mouse. Striatal inhibitory neurons (Cluster 06 Inh_striatal) expressed inhibitory markers *GAD2* and lower-layer marker *BCL11B* [36]. Layer 19-P (Cluster 09 layer[19P]) expressed canonical Cajal-Retzius cell markers *RELN* and Purkinje cell marker *PCP4*. In addition, they also expressed non-coding RNAs *SNHG11* and *MEG3* (Fig. 2D, S2C). Microglia (Cluster 16 Microglia) had higher cell numbers in human than in mouse and exhibited low expression correlation with other clusters. They expressed microglial markers such as *LAPTM5, C1QA, C1QB*, and *C1QC* [37], as well as several microglia-specific pathways such as TREM2-TYROBP pathway [37-40] (Fig. 2D). Interestingly, Cluster 20 Unknown exhibited an intermediate state between excitatory neuron and interneuron: it had high expression correlation with both excitatory neuron and interneuron, and expressed several marker genes such as *STMN2* (excitatory neuron) and *NNAT* (interneuron) (Fig. 2D, S2C, D).

### Different developmental trajectories of excitatory neuron between human and mouse

To compare the developmental trajectories in excitatory neuron between human and mouse, we split the data into 6,623 human cells and 6,655 mouse cells, and performed unbiased clustering using UMAP again. We identified 23 clusters and also observed lineage separation of the major cell types (Fig. S3A, B, C, D). We used *NEUROD6* as marker gene and identified 6 excitatory neuron clusters in human (Cluster 00 SVZ, 01 Exc deep, 02 Exc deep, 03 Exc upper, 04 Exc upper and 05 Exc upper) and 7 excitatory neuron clusters in mouse (Cluster 00 SVZ, 01 SVZ, 02 Exc deep, 03 Exc deep, 04 Exc deep, 05 Exc upper and 06 Exc upper) (Fig. S3A, B). We then reconstructed the corticogenesis developmental trajectories using Monocle analysis. Both human cells and mouse cells were distributed along pseudo-temporally ordered paths from SVZ to neocortical layers, including deeper neocortical layers (layer VI, then layer V) and superficial layers (layer IV, then layer II/III) [28] (Fig. 3A, B). Surprisingly, we found that the developmental trajectory in mouse was a continuous process without any branch, in which SVZ migrated to deeper neocortical layers and then to superficial layers. By contrast, in human this process had two branches in which SVZ migrated to both deeper neocortical layers and superficial layers (Fig. 3A, B, S3E, F). Consistent with this observation, we found that SVZ marker *EOMES* was specifically expressed in the beginning of trajectories, deep-layer marker *BCL11B* was specifically expressed in the upper branch in human, whereas migratory and upper-layer markers *POU3F2* and *SATB2* were specifically expressed in the lower branch, which was different from those in mouse trajectory (Fig. 3A, B, S3I). More important, when we further split the data based on development stages, we still found that the lower-layer neurons were present in 7 to 23 pcw in human and E14.5 and P0 in mouse, with high expression of *BCL11B*. Meanwhile, a large number of Layer II-IV (upper layer) cells at P0 in mouse and 20-23 pcw in human was identified, with high expression of *SATB2* (Fig. S3G, H).

**Figure 3.**
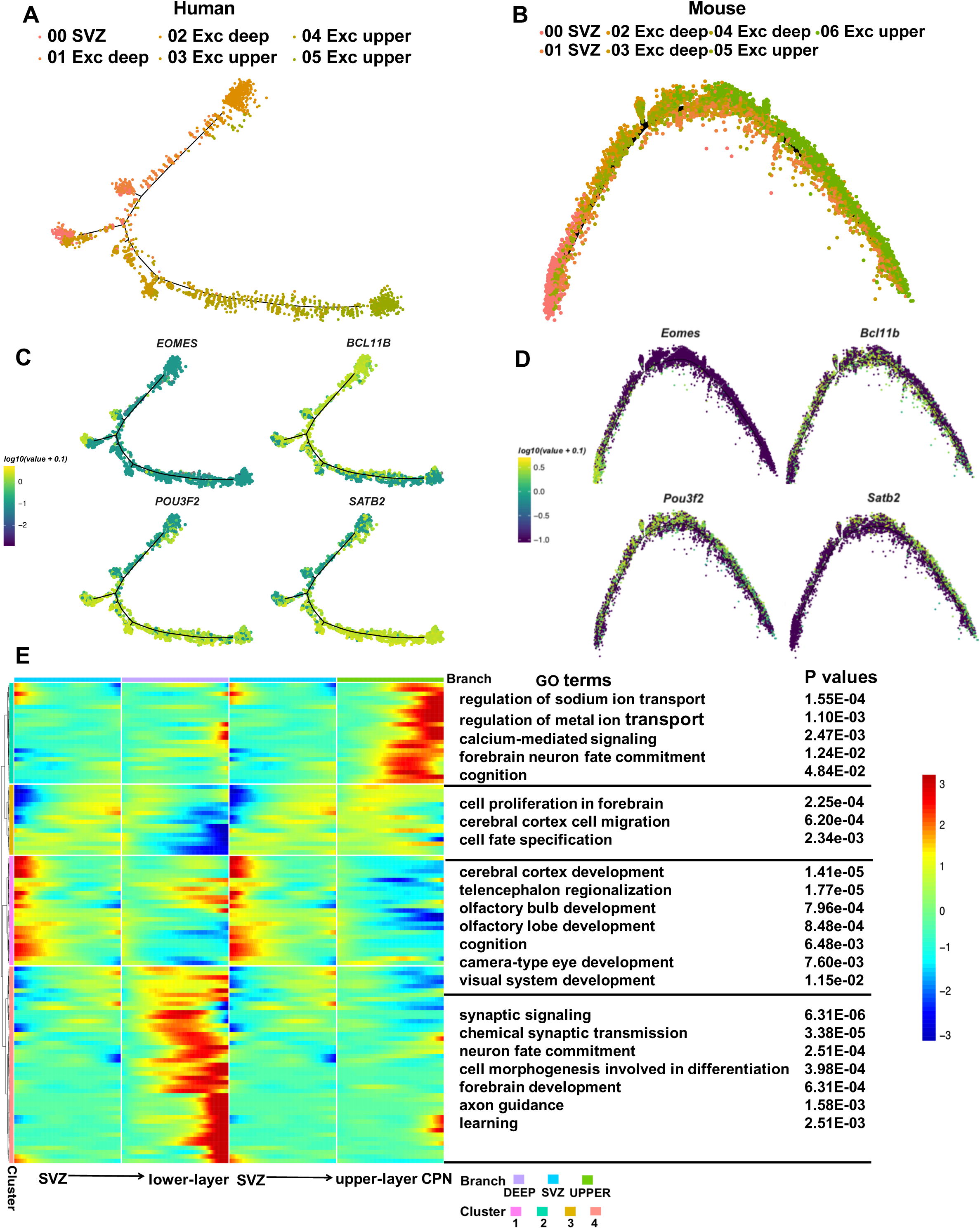

We identified 109 differentially expressed genes in human that may determine cell fate commitment. They were fall into 4 clusters which exhibited successive waves of gene expression in different branches (Fig. 3E). In addition, Gene Ontology (GO) enrichment analysis revealed that the functions of genes predominantly expressed at the SVZ branch were enriched in cerebral cortex development, olfactory bulb development and olfactory lobe development as previously reported [41]. Among these genes, transcription factor *PAX6* is crucial for SVZ neurogenesis and also acts via retinoid signaling to regulate eye development [42] suggesting that eye development may have relative to SVZ neurogenesis (Fig. 3E). Lower-layer from human cortex showed increased expression of genes for modulation of synaptic signaling and chemical synaptic transmission, which have a mechanistic link with learning and memory by plasticity regulation [43]; whereas upper layer CPN from human cortex exhibited increased expression of genes involved in regulation of sodium ion transport and calcium-mediated signaling (Fig. 3E). Genes related to forebrain neuron fate commitment and forebrain neuron differentiation were expressed in both lower-layer and upper layer. More interestingly, the functions of genes that were highly expressed in SVZ, lower layer and upper layer were all enriched in cognition and learning (Fig. 3E).

We further analyzed the migratory marker genes expression in neocortical. We found these genes had continuously increased expression in SVZ, and decreased expression in deep-layer and upper-layer in human. Specifically, *NRP1* and *PALMD*, which were involved in the regulation of cell morphogenesis and cerebral cortex cell migration, exhibited increased expression in the SVZ lineage and then declined in deep layer and upper layer lineage, suggesting that neuronal migration in projection neurons in the cerebral cortex may occur from the SVZ’s maturity stage to the early development of deep and shallow layers, rather than the late deep layer [44] (Fig. S3J, K). By contrast, in mouse cerebral cortex we found migratory genes showed increased expression in SVZ and deep layer, and decreased expression in upper layer suggesting that in mouse the deep layer continuously process cell migration from SVZ to upper layer. Representative genes include *Meis2, Nrp1* and *Pou3f2*, whose functions are relative to cell migration and regulation of cell morphogenesis involved in differentiation (Fig. S3K).

### Divergence in neocortical between human and mouse

In order to better understand the distinct layers in neocortical in human and mouse, we performed a comparison of cortical expression models through single-cell analysis, including deeper neocortical cortex (layer VI, then layer V) and late-born neurons migrate through deep layers to gradually fill more shallow layers (layer IV, then layer II/III). Pairwise comparison of human and mouse cortical layer data sets identified a large number of differentially expressed genes (DEGs) (Fig. 4A). We observed a considerably bigger number of differentially expressed genes in human than in mouse, meanwhile we also found some co-expression marker genes, for example, fez family zinc finger 2 *Fezf2* (high expressed in deeper layers), special AT-rich sequence binding protein 2 (*SATB2*) and *POUEF2*, a marker of layer II/III with function of migration and differentiation [28]. More interestingly, we detected the differentially expressed gene: *CLSTN2*, high expressed in human deep layer and mouse upper layer, which significantly associated with pathway of cognitive and memory behavior and reduction in functional inhibitory synapses [45], suggesting that genes, relating to cognition and memory pathways, expressed earlier in human than mouse.

**Figure 4.**
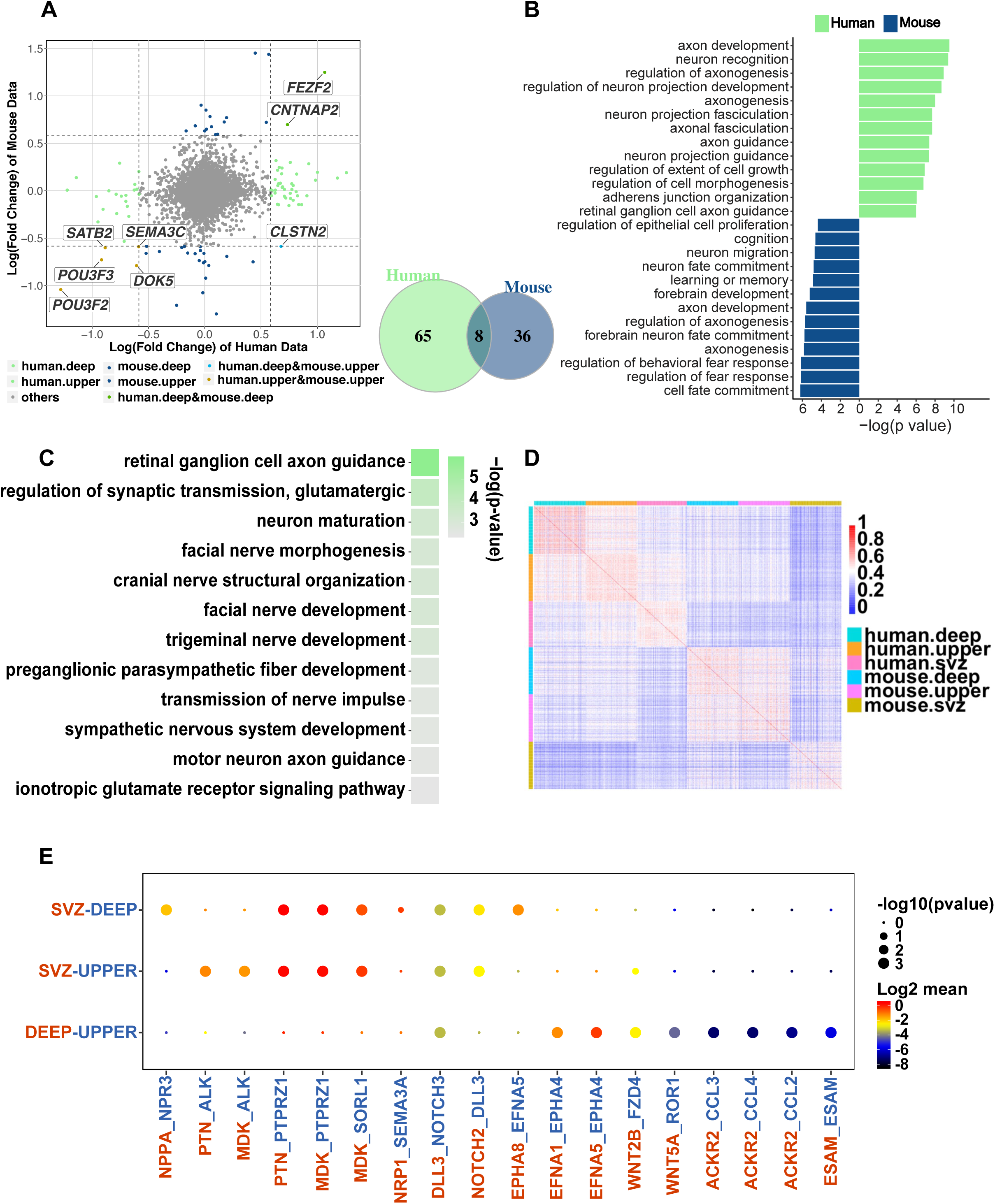

Pathway analysis of two layers specific to humans revealed enrichment of axon development, such as axon guidance and axonal fasciculation, compared with mouse (Fig. 4B). Additionally, enriched pathway contained in many aspects, including visual system, facial nerve, trigeminal nerve and motor neuron in human — more specifically, retinal ganglion cell axon guidance, facial nerve development, trigeminal nerve development, motor neuron axon guidance and regulation of synaptic transmission, glutamatergic (Fig. 4C). In contrast, some pathways were enriched in mouse, mainly in cell fate commitment, neuron fate commitment and neuron migration (Fig. 4C).

To continuously process research in neocortical in human and mouse, we expanded our analysis included SVZ. Pairwise comparison of human and mouse SVZ and upper cortical layers scRNA-seq transcriptomics data identified a large number of DEGs (Fig. S4A). The number of DEGs detected in mouse was 6 times higher than that derived from human. SVZ marker genes, *EOMES* and *NHLH1*, related to cell-type determination within developing neurons, upper layer markers, *SATB2* and *MEF2C*, had co-expression in human and mouse. We found more than 200 significant gene ontology (GO) terms in human and mouse respectively. Specifically, for human, pathways focused on neuron migration and axon development. In addition, human-specific genes also revealed enrichment of “eye development”, “visual system development”, and “sensory system development”. For mouse, we detected mouse-specific pathways “learning or memory”, and “synapse organization” (Fig. S4B). we also identified the DEGs between SVZ and deeper layer among human and mouse (Fig. S4C). The number of DEGs are equal in human and mouse (Fig. S4D).

We next examined the correlation between each cluster in neocortical between human and mouse (Fig. 4D). We identified mouse SVZ and mouse upper layer showed remarkably low degree correlation (Fig. 4D). In contrast, the correlation of human SVZ with human deep layer were nearly similar to those of human SVZ with human upper layer (Fig. 4D). Taken together, we conclude that the differences of deeper layer and upper layer has larger in human than mouse, while the differences of SVZ and upper layer has larger in mouse than human (Fig. 4D).

Having defined the populations of SVZ, deeper layer and upper layer cells, we further used CellPhoneDB to perform an unbiased ligand receptor interaction analysis between these populations [46]. To detect how neocortical projection neurons are generated in human, we focused functional analyses on interactions with SVZ, deeper layer and upper layer (Fig. 4E). SVZ expressed *PTN* and *MDK*, signaling to receptors *ALK* on upper layer, that are known to regular nervous system development, function, and repair, especially, regulating the sympathetic neuron growth during development and aberrant signaling to neuroblastoma predisposition [47-49] (Fig. 4E). SVZ expressed *PTN* and *MDK*, signaling to the receptor tyrosine phosphatase family *PTPRZ1* on deeper and upper layers, which shows the significant function in activate cell growth, migration and cellular activities (Fig. 4E). More importantly, *PTPRZ1* interaction with *MDK* promotes neuron migration [50]. SVZ expressed high levels of *NOTCH3* interacting with receptor *DLL3* on deeper layer cells and upper layer cells, which could modulate cellular differentiation, fates for terminal neuronal differentiation and links to progression toward a neuronal fate in the developing cortex [51, 52](Fig. 4E). In addition, ephrin family showed high level of expression in human cortical. Firstly, *EPHA8_EFNA5* could promotes neurite outgrowth and axon guidance [53], which showed significance in process SVZ to deep layer (Fig. 4E). *EFNA1_EPHA4* involves in dendritic spine morphogenesis in deep layer to upper layer [54] (Fig. 4E). Deep layer expressed *WNT* genes interacting with receptors *FZD4* and *ROR1* on upper-layer cells. *WNT5A_ROR1* promotes the formation of presynaptic [55] (Fig. 4E). Upper layer expressed Atypical Chemokine Receptor 2 ligands (*ACKR2*) receptors that are known to be required for immature/mature dendritic cells [56] (Fig. 4E). *ESAM_ESAM* helps cell-cell adhesion in deep layer to upper layer (Fig. 4E).

In summary, our trajectory, cell-type and unbiased ligand-receptor interactions analysis between mouse and human indicates that human and mouse have differences in development neocortical models, the correlation between deep layer and upper layer are lower in human than mouse, and the ligand-receptor analysis indicate that deep layer and upper layer may not have various neuron migration interactions in human. These phenomena may therefore show SVZ migration to appropriate layers are relatively independent in human neocortical projection neurons.

### Characterization of the excitatory neuron gene expression pattern across species

To better understand the divergence and conservation of excitatory neuron gene expression across evolution, we used developmental time course reconstructed by monocle analysis to process the further analysis of gene expression pattern. Although a large part of genes keep conservation across evolution, some genes showed converse expression trends in human and mouse. We defined two gene expression clusters, with cluster a representing up-regulated genes during the frontal development excitatory neuron in human, mainly including SVZ cells (Fig. 5A), and in mouse, the expression patterns are converse (Fig. 5B). Cluster a contained genes *VCAN, BCL11B, SOX5, PTN* and *CNR1*, whereas Cluster b contained genes *ST18, KLHL13* and *NHLH2* (Fig. 5A, B). We focused some genes have key functions in neuron development, for example, we detected *CNR1* have various difference in expression pattern, in human, it expressed in SVZ cells, whereas it expressed in deeper and upper layer cells in mouse, in other words, CNR1 expressed earlier in human than mouse in cortex (Fig. 5A, B, C). CNR1, known as cannabinoid receptor 1, is expressed in the peripheral nervous system and central nervous system and combine cannabinoid to regulate the release either glutamate or GABA, and the endocannabinoids (*eCBs*) and their receptors (*CNR1*) express in cortex that can act as the significant function as cell proliferation and migration and axon guidance during the critical stages of brain development [57-62]. In addition, *CNR1* also likely plays an important role in the development of memory and cognition, and is relative to many emotional, neurological diseases, such as novelty [63-65]. *ST18*, expressing earlier in mouse than human, shows key factor in breast cancer [66] (Fig. 5A, B, D).

**Figure 5.**
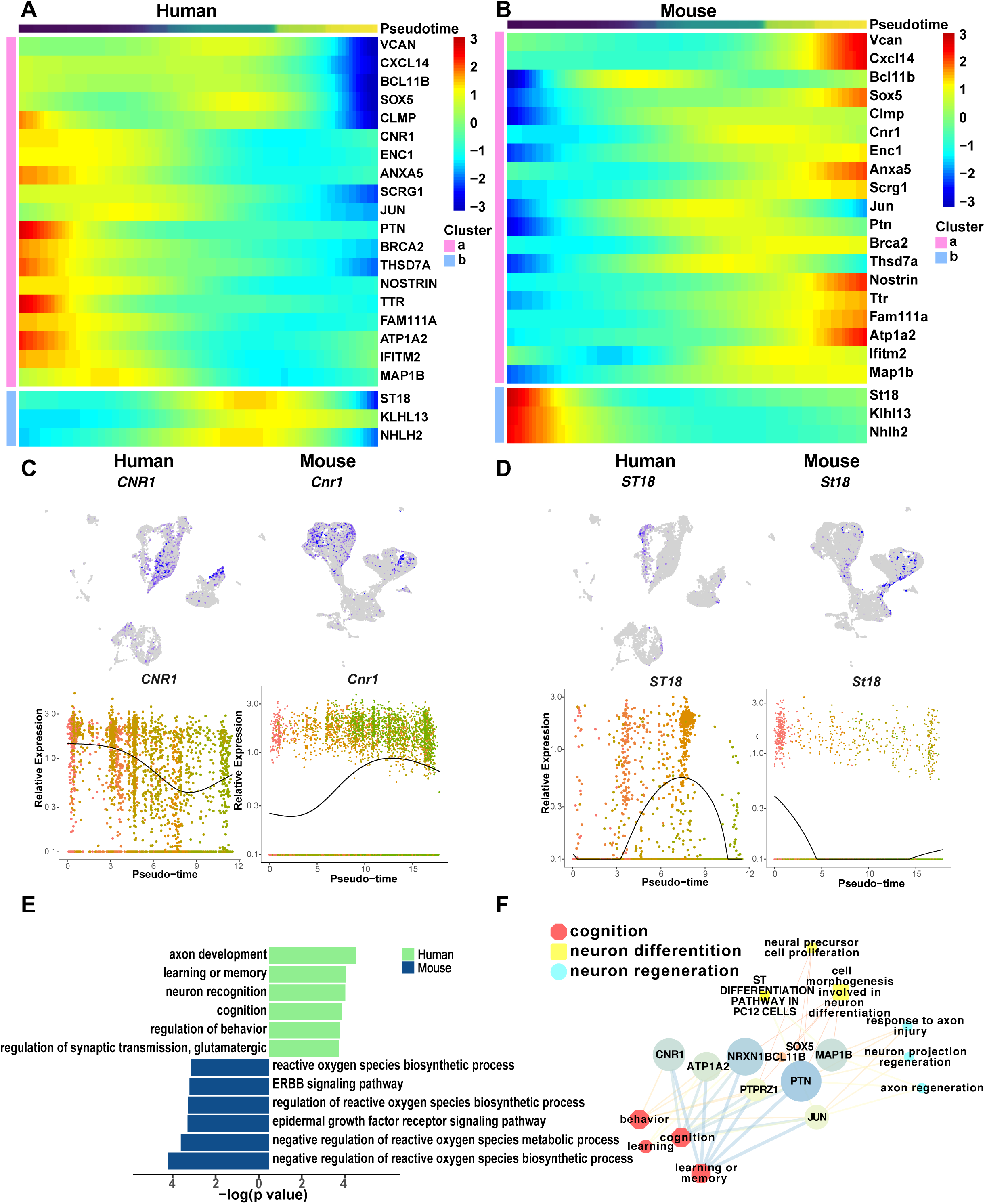

In order to detect the function of cluster a genes and cluster b genes, we found more genes with similar expression patterns to process GO terms, function protein network and transcriptional regulatory relationships analysis. We detected some neuron differentiation and synaptic pathways, including chemical synaptic transmission, cell morphogenesis involved in neuron differentiation, active in human SVZ (Fig. S5E, F). More interesting, axon development, learning or memory, neuron recognition, cognition, regulation of behavior and regulation of synaptic transmission, glutamatergic were more active in the human SVZ than mouse SVZ (Fig. 5E). The key genes, enriching in cognition, learning or memory, neuron differentiation and neuron regeneration pathways, including *PTPRZ1, PTN, CNR1, ATP1A2, SOX5* and *JUN*, showed the different expression pattern in human and mouse excitatory neuron, suggesting human probably develop these function, especially for cognition, learning or memory, earlier than mouse (Fig. 5F, S5A, B, C, D). According to protein and transcription factor analysis, we observed key protein or transcription factor: *JUN* connected with *FOS*, as a classic marker for neuronal activation, which has important regulator of axonal regeneration and neuronal activation. [67] (Fig. 5A, 5B, S5E, F). Moreover, *JUN-PTN* was detected TF-target regulatory relationship, which directly indicated activation of neuronal migration during later brain development [68-70], and ligand-receptor interaction: activating *PTN_PTPRZ1* in human SVZ to deeper layer and SVZ to upper layer before (Fig. 4E), which activates cell growth, migration and cellular activities [70, 71], and we found these genes have different expression pattern in human and mouse, for human, they showed highly expression in SVZ, whereas *Ptprz1* and *Ptn* performed low expression in mouse SVZ, and increase expression in deeper layer and upper layer cells (Fig. 5A, B, S5A, B), indicating that neuron cell migration performs in human SVZ to deeper layer and upper layer or mouse deeper layer to upper layer process. We also found genes expressing in mouse SVZ, like *ST18* and *UACA*, relative to reactive oxygen species biosynthetic process (Fig. 5D, E).

### Single-cell transcriptomic analysis identifies human excitatory neuron subsets

In order to examine whether the excitatory neuron of human and mouse display similar cell types or contain several subtypes, we further observed the clustering of human and mouse (Fig. S3A, B). Surprisingly, distinct clusters of maturing neurons (04 Exc upper and 05 Exc upper) reproducibly emerge according to the distinct expression profiles, including reciprocal expression of PFC (The prefrontal cortex) marker (*ABCA1*) and V1 (Visual cortex) marker (*LPL*) [9] (Fig. 6A). By contrast, the gene with reciprocal expression in human 04 Exc upper and 05 Exc upper don’t have similar expression patterns in mouse (Fig. 6B).

**Figure 6.**
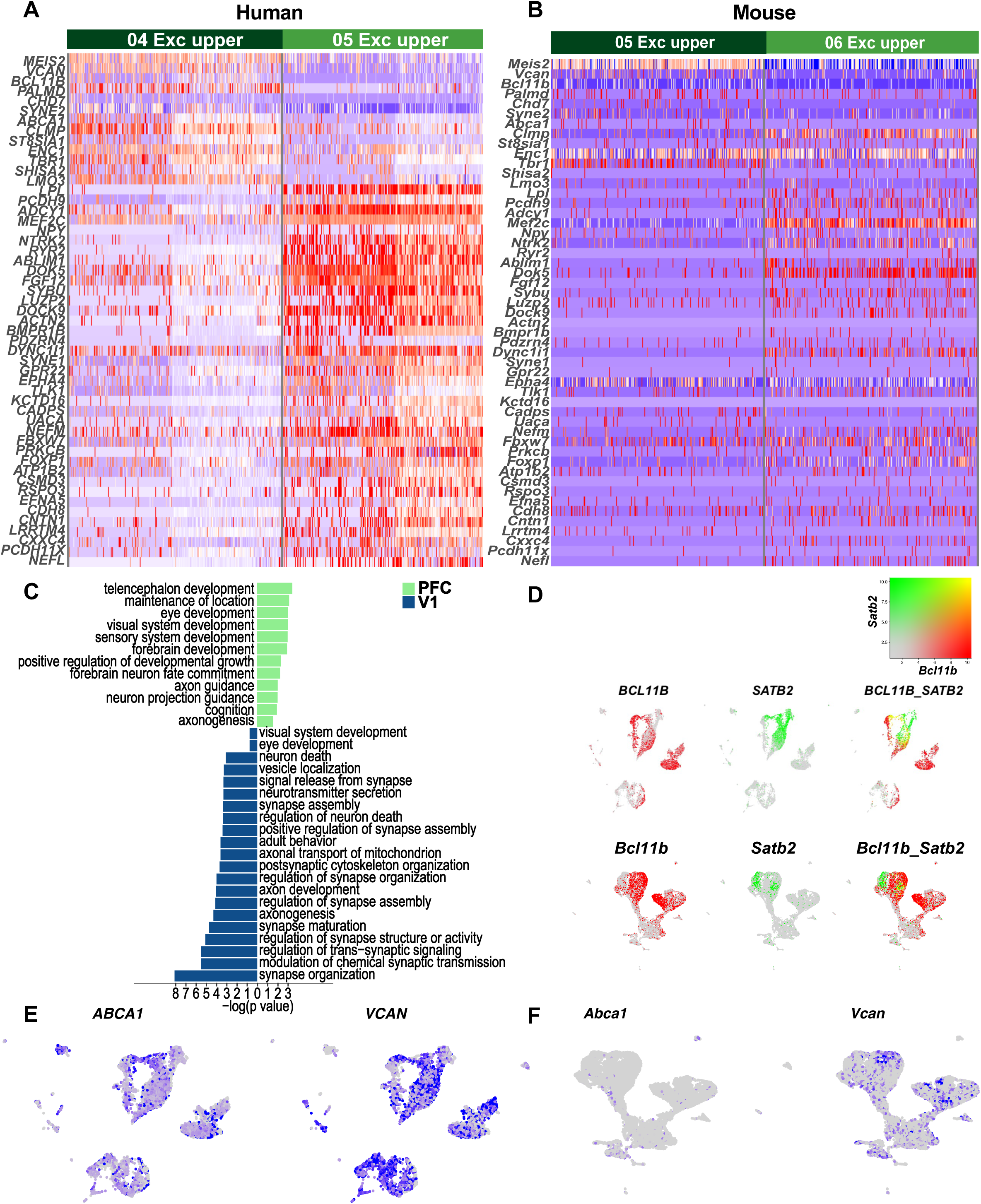

In order to further make sure the cell types, we used known labels to tag the clustering and processed statistics (Fig. S6A, B, C, D), we found that the proportion of PFC and V1 cells have remarkable difference in 05 Exc upper (Fig. S6C), suggesting that the coarse laminar distinctions among newborn neurons are resolved into refined transcriptional state in human later development, whereas the phenomenon disappeared in cross-species clustering that the number of PFC and V1 cells in each cluster are similar, especially in excitatory neuron, indicating that the newborn neurons probably don’t have refined expression states in mouse among the E14.5 and P0 stages (Fig. S6D).

In order to detect the biological processing of PFC and V1 in human, We therefore selected expression differentially genes for pathway analysis by gene ontology resource and functional protein association networks (STRING) [72](Fig. 6C, S6D, E, F). Axon guidance, axonogenesis, axon development were active in PFC and V1 (Fig. 6C). Pathway relative to synapse signaling, including synapse organization, modulation of chemical synaptic transmission, regulation of trans-synaptic signaling and synapse structure or activity, synapse maturation, axonal transport of mitochondrion, were more active in V1 (Fig. 6C). In PFC area, we enriched sensory system development and cognition, rather in V1, and in PFC and V1, we all discovered visual system development and eye development (Fig. 6C).

To further examine the notion of typological distinction between PFC and V1 inner human and mouse, we compared distinct co-expression profiles of genes involved in the specification of projection patterns during development, for example, *SATB2* and *BCL11B* co-expressed in human maturing PFC deep layer and upper layer at later stages of development (Fig. 6D). In contrast, neurons in V1 had distinguish expression pattern of *BCL11B* and *SATB2* (Fig. 6D). More interestingly, *Bcl11b* and *Satb2* also had co-expression phenomenon in mouse, but it appeared in the immature neurons, indicating that the early fate specification [18] (Fig. 6D). Then, we found that it is common that maturing PFC neurons co-expressed genes in deeper and upper layer cells, for example, *ABCA1* and *VCAN*, which couldn’t express in deep layer in mouse (Fig. 6E, F). Additionally, we focused on the function of these genes, which may play role in intercellular signaling and extracellular structure organization.

### Conservation of radial glia developmental trajectories

The comparison of excitatory neurons is crucial for our understanding of many differences between human and mouse developing cortex. However, they are many cell types that haven’t been compared. We further analyzed the developmental trajectories in radial glia. We observed there are two RG clusters in cross-species clustering (cluster 10 RG and 11 RG) (Fig. 1D), which had similar human and mouse cells proportions (Fig. 2A, B). During early development, dorsal telencephalic radial glia produces excitatory cortical neurons, while ventral telencephalic produces inhibitory cortical interneuron during 7 to 23 pcw in human that have been reported [9]. We applied principal component analysis (PCA) for cross-species clustering that observed 11 RG migrate to 13 GE (Interneuron migrate tangentially from the ganglionic eminences) and 12 G2/M, and 10 RG migrate to 00 SVZ (Fig. S7A, B), and then, we chose these clusters to reconstruct the trajectories analysis (Fig. 1D), that we found two lineages: from 11 RG to 13 GE and 12 G2/M or 10 RG to 00 SVZ (Fig. 7A). Group of cells expressing *MKI67*, markers of proliferating and glia, were detected at early timepoint at the pseudo-time analysis by Monocle, while *EOMES* and *NEUROD6* expressed in SVZ branch, and *GAD2* expressed in GE branch (Fig. 7B). We detected that these developing processes have been emerged at stage 2 (E14.5 in mouse and 7-to 11.5 pcw in human). We use species-specific clusters to analysis with four markers to depict the expression profile (Fig. S7B, C, D, E).

**Figure 7.**
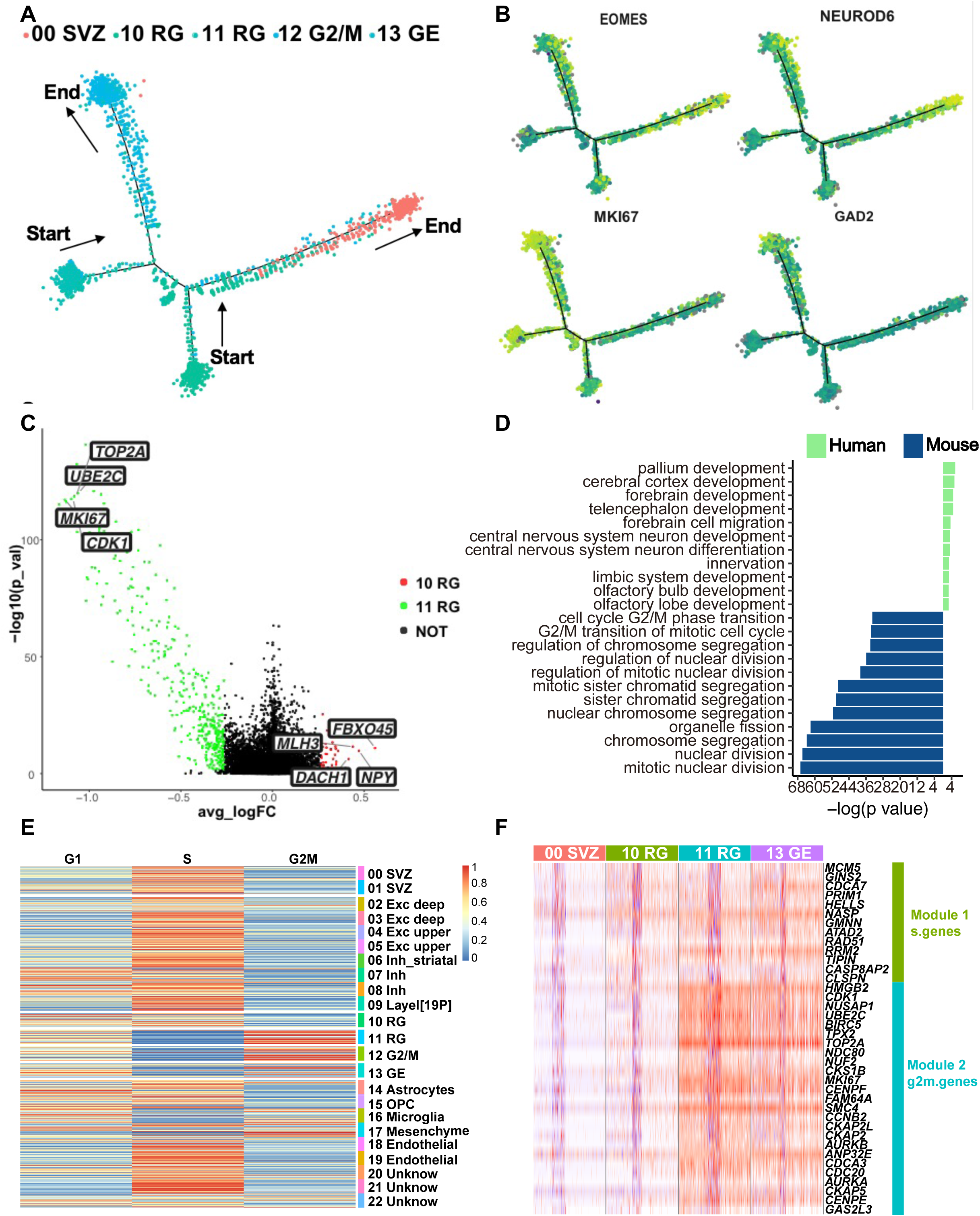

In order to detect the gene expression relative to cell fate commitment, we identified 80 genes highly dispersed in human and mouse. DE genes that fall into 5 clusters and depicted successive waves of gene expression in different branches (Fig. S7F). Genes predominantly expressing at the 11 RG branch enriched in pathways, including nuclear division, mitotic nuclear division and sister chromatid segregation, suggesting that these cells stayed in G2/M stages (Fig. S7F). Genes co-expressed in 10 RG and 11 RG enriched in epithelial cell proliferation, gliogenesis, astrocyte differentiation, as expected in radial glia (Fig. S7F). GE cells expressed genes, relating to cerebral cortex GABAergic interneuron differentiation (Fig. S7F).

To examined two RG expression profile and enrichment pathways in human and mouse, we detect expression differentially genes in these two clusters (Fig. 7C). 11 RG expressed some proliferating marker genes, including *MKI67, TOP2A* and *UBE2C*, and regulate cell cycle genes, like *CDK1* and *TPX2*, which co-expressed in human and mouse ventral telencephalic radial glia (Fig. 7C); while dorsal telencephalic radial glia expressed three kinds genes, *HES5* (RG marker), *EOMES* (SVZ markers) and *DACH1* (enriched in ventricular radial glia cells) [73] (Fig. 7C), that related to olfactory bulb development and olfactory lobe development, which also enriched in SVZ (Fig. 7D).

As many proliferating and cell cycle genes high expressed in 11 RG, rather than 10 RG, we further analysis the cell cycle of cell types. Excitatory neuron exhibited a profile highly similar to interneuron that stay at S states, and 10 RG kept at G1/S states. In contrast, 11 RG, 12 G2/M and 13 GE continued cell division (Fig. 7E, F).

## DISCUSSION

We have systematically analyzed scRNA-seq data to integrate and compare developing human and mouse cerebral cortex, including identified cell type diversities, developing trajectories, gene expression patterns, functional protein association networks, signal transduction pathways and ligand receptor interaction analysis.

According to previous reports, neocortical projection neurons are generated in an “inside-out” fashion by diverse progenitor types in the VZ and SVZ, for example, cut-like homeobox 2-positive (*Cux2+*) progenitors produce the most superficial-layer (upper) [74, 75], which is resided callosal projection neurons (CPN) and high expression marker gene (*SATB2*), whereas corticofugal-layer (deep) neurons derive from Cux2-negative (*Cux2–*) progenitors, which include a large number of corticothalamic projection neurons (CThPN) that reside in layer VI, and subcerebral projection neurons (SCPN) that reside in layer V and express *BCL11B* and *FEZF2* [28]. As the result of the basis of current evidence, a number of different models are produced, In line with a previous report, we detected the ‘sequential competence states’ model is undesirable in mouse and human [28], because the mouse early developing deep layer (02 EXC deep) expressed an upper layer marker (*SATB2*), a deep layer marker (*BCL11B*), a migratory marker (*TIMA2*) and a transcription factor that is expressed in Layer II–III neurons during their migration and differentiation (*POU3F2*) (Fig. S8B), and human developing trajectories (Fig. 3A), suggesting that there is unreasonable that individual progenitors (*Cux2–*) produce a single neuronal subtype: CThPN at a time [28] (Fig. S8F). More significantly, according to our study, we indicated human and mouse exist two developing models. For human, we observed the reconstructed excitatory neuron trajectory, indicating that the deep and upper layer migrate from SVZ independently (Fig. 3A), whereas the deep layer and upper layer ordering are chaos in mouse developing neocortex trajectory (Fig. 3B), and then, we detected the ligand and receptors have differentially expression in human and mouse (Fig. 4E, S5A, B).

Additionally, *CUX2* expressed differentially in human and mouse, for example, *CUX2* only expressed in upper layer in human (Fig. S8C), while *Cux2* have expression in maturing mouse deep layer neuron (Fig. S8D), indicating that *Cux2-* positive lineage after generating deep-layer neuronal sub-types change their competence state to express *Cux2* and produce CPN to generate upper-layer. The results may suggest that human is more likely to similar the model “independent lineages” and the model “nested multipotential lineages” is more likely to exist in mouse [28] (Fig. 8A).

**Figure 8.**
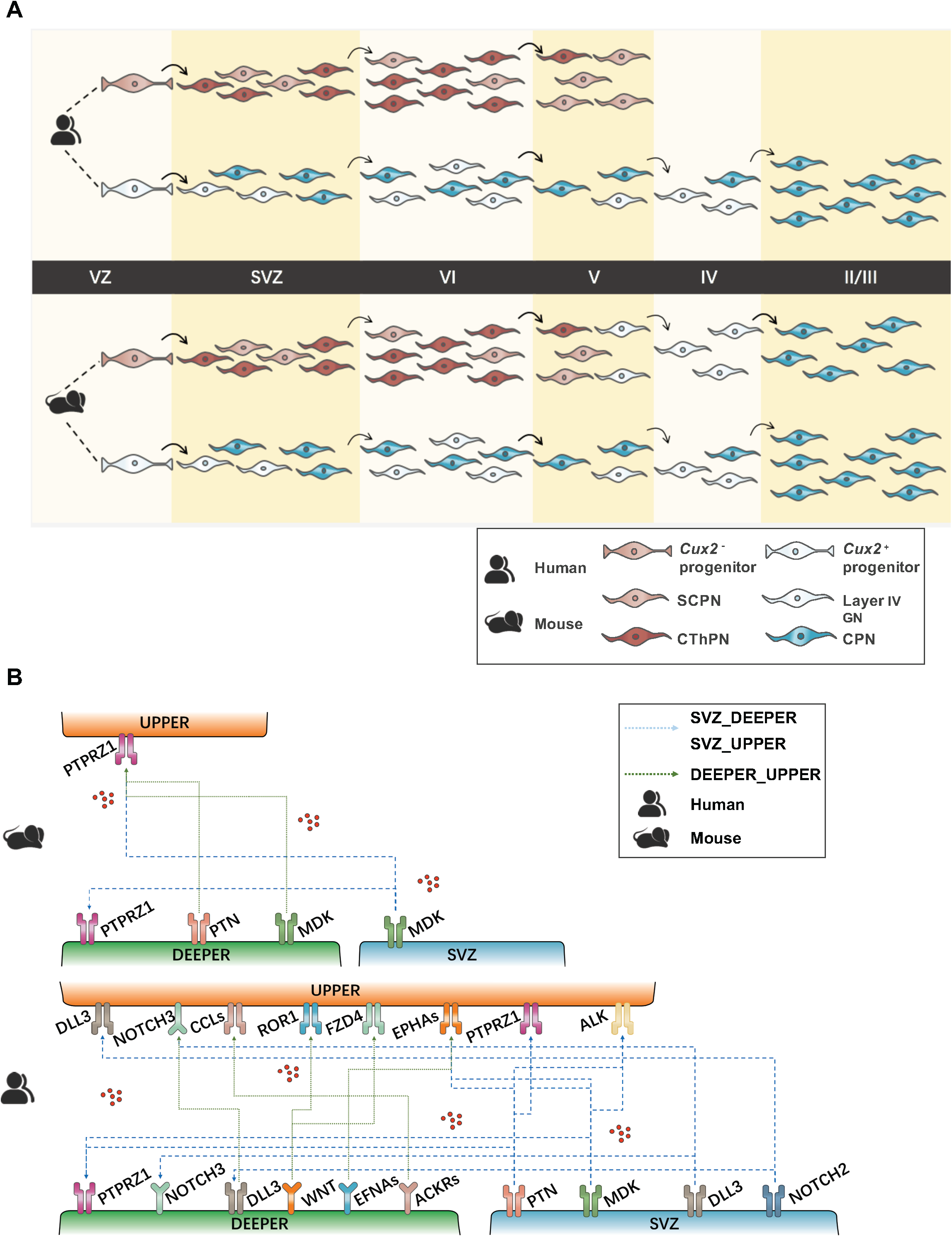

We further summary the divergence and conservation of ligand receptor interaction analysis between human and mouse neocortical projection neurons [46] (Fig. 8B). We predicted the neuron migration and differentiation between SVZ and layers that could potentially be triggered by protein-interactions: *PTN_PTPRZ1* and *MDK_PTPRZ1* [50] (Fig. 8B), meanwhile these expression patterns show high expressed in human SVZ to deep layers and upper layers, and mouse SVZ to deep layers and upper layers, and deep layers to upper layers, which are consistent with our predicted two models: human for “independent lineages” and mouse for “nested multipotential lineages” (Fig. 8B, S8E, F). Then, we observed *NOTCH, EFNA, EPHA, WNT, ACKR* and *CCL* signaling in human deep and upper layer with significant function in the development of synaptic and dendritic cells [53] (Fig. 8B).

Our data also suggest that cognition-linked genes might form robust groupings according species-specific and cell-type expression profiles. We detected many cognition-relative genes expressed earlier in human than mouse, including *PTPRZ1, CNR1, PTN*, and *CLSTN2* (Fig. 4A, 5A, B, C, S5A, B), that enriched in cognition, learning and memory and behavior pathways [45, 63, 76] (Fig. 5E, S5F), therefore understanding human cognition development is probably earlier than those of mouse. In conclusion, our single-cell analysis provides an essential resource for future studies directed at understanding the difference developing neocortical trajectories, and excitatory neuron sub-types expression profiles, and cognition-relative genes expression profiles during the early human and mouse brain development.

## METHODS

### Data processing of scRNA-seq from Chromium system

In the early data processing, we applied the processed gene-cell data matrix without poor-quality cells from published study. We used Seurat package (v3.1.2) to process secondary filtration [77]. Only cells that expressed more than 500 genes were considered, only genes expressed in at least 5 single cells were included for further analysis. Additionally, we selected the homologous genes between human and mouse to process further analysis. In total, 12350 genes across 18446 mouse and 6623 human single cells remained for subsequent analysis. The data were normalized to a total of 1 × 10^4^ molecules per cell for the sequencing depth using the Seurat package. The batch effect was mitigated by using the ScaleData function of Seurat (v3.1.2).

### Data integration of scRNA-seq and identification of cell types

The Seurat package (v3.1.2) was used to integrate human and mouse scRNA-seq data. We applied the new normalization method ‘SCTransform’ to process cross-species clustering in order to better operation downstream integration [77]. Then, the Seurat package was used to perform linear dimensional reduction. we used SelectIntegrationFeatures or FindVariableFeatures function from Seurat and further analysis used ‘FindIntegrationAnchors’ to select 2000 highly variable genes as input for PCA. Then we identified significant PCs based on the JackStrawPlot function. Strong PC1-PC30 were used for UMAP to cluster the cells by FindClusters function with resolution 0.8. Clusters were identified by the expression of known cell-type markers. The markers EOMES, NEUROD6, GAD2 and MKI67 were used to identify the basic cell-type, including progenitor cells, excitatory neurons, inhibitory neurons and astrocytes, respectively.

### Identification of DEGs between clusters

The DEGs of each cluster were identified using FindAllMarkers function (min.pct = 0, logfc.threshold = 0, test.use = “wilcox”) with the Seurat R package [77]. We used the Wilcoxon rank-sum test (default), and top 10 DE genes (p < 0.05) by fold change were identified for each cluster.

### Correlation Analysis

For correlation analysis of merged human single cell and mouse single cell data, we used integrated data from Seurat package and the Pearson-correlation coefficient calculated in Hmisc package (version 4.3-0), and the relative visualization used package pheatmap (version 1.0.12).

### Constructing single cell trajectories in cerebral cortex

The Monocle 2 R package (version 2.10.1) were applied to construct single cell pseudo-time trajectories to discover developmental transitions [78]. We used data matrix from Seurat with 2000 highly variable genes into pseudo-time order. we then called ‘differentialGeneTest’ to process DEGs analysis in different cell-types. ‘DDRTree’ were applied to reduce dimensional space and “orderCells” were used to sort cells. The minimum spanning tree on cells and marker genes expression profiles were plotted using the visualization functions “plot_cell_trajectory” for Monocle 2

### Identifying genes relative to cell fate choices

We analyzed branches in single cell trajectories by BEAM and calILRs function, and we selected genes with q-value < 0.01 and a mean expression value ≥ 0.18, Genes dynamically expressed along the different branches were clustered using the plot_multiple_branches_heatmap function and plot_genes_branched_heatmap function, and Enriched GO terms of cluster-grouped genes were identified using clusterProfiler package (version 3.10.1) [79]. Then, genes were clustered into different clusters by using ‘plot_multiple_branches_pseudotime’ and ‘plot_genes_branched_pseudotime’ to visualize the expression profiles in different cell-types.

### GO enrichment analysis

Gene ontology (GO) biological process enrichment analysis were performed on differently expressed genes (DEGs) (p-value < 0.05, fold change (FC) > 1) using clusterProfiler package (version 3.10.1) and Metascape [79, 80].

### Analysing inter-lineage interactions within the neocortex

For comprehensive systematic analysis of inter-lineage interactions within the neocortex, we use CellPhoneDB [46]. We used gene-cell matrix data from Seurat packages after integration as the input data, and we considered only ligands and receptors expressed in greater than 10% of the cells in any given subpopulation. Then, we seted the P-Value 0.05 to ensure the prominence. We called ‘cellphonedb plot dot_plot’ to visualize the ligand-receptor interactions within the neocortical projection neuron.

### Comparing gene expression pattern divergence

The pseudo-time of each cell was assigned by Monocle2, and used median function in R to calculate the mean of pseudo-time in mouse and human trajectories, dividing cells into two categories, including early stage of the cells and the latter stage of cells. and we reconstructed the clustering included 132500 homologous genes within human and mouse by using Seurat package [77, 78]. FindMarkers function helps us to calculate the expression profiles between these two groups, and genes with average expression difference > 0.585 natural log with *P* < 0.05 were selected as DEGs between human and mouse. a part of genes visualized by ‘plot_pseudotime_heatmap’ function.

### Functional protein association networks analysis

Functional protein association networks were calculated on DEGs using STRING, the results visualized using Cytoscape [72, 81].

### Connecting transcription factors to target genes

To find the potential transcription factors, TRRUST was used for search transcriptional regulatory interactions [82], and the results visulized using Cytoscape [72, 81].

### Cell-cycle analysis

In the cell-cycle analysis, we applied scran packge in R (version 1.10.2) to score G1, S and G2/M states of each cells [83]. we selected a cell-cycle related gene set with 43 genes expressed during S and 54 genes expressed during G2/M from Seurat package [77]. These gene sets should be anti-correlated in their expression levels, and cells expressing neither are likely to be in the G1 phase (not cycling).

### Statistical analysis

Comparisons between two groups were made using wilcox. Sample size and P values are given in the Figure legends.

## ACKNOWLEDGMENTS

This work was supported by the Strategic Priority Research Program of the Chinese Academy of Sciences (Grant No. XDA16010114)

---

**Figure S1.**
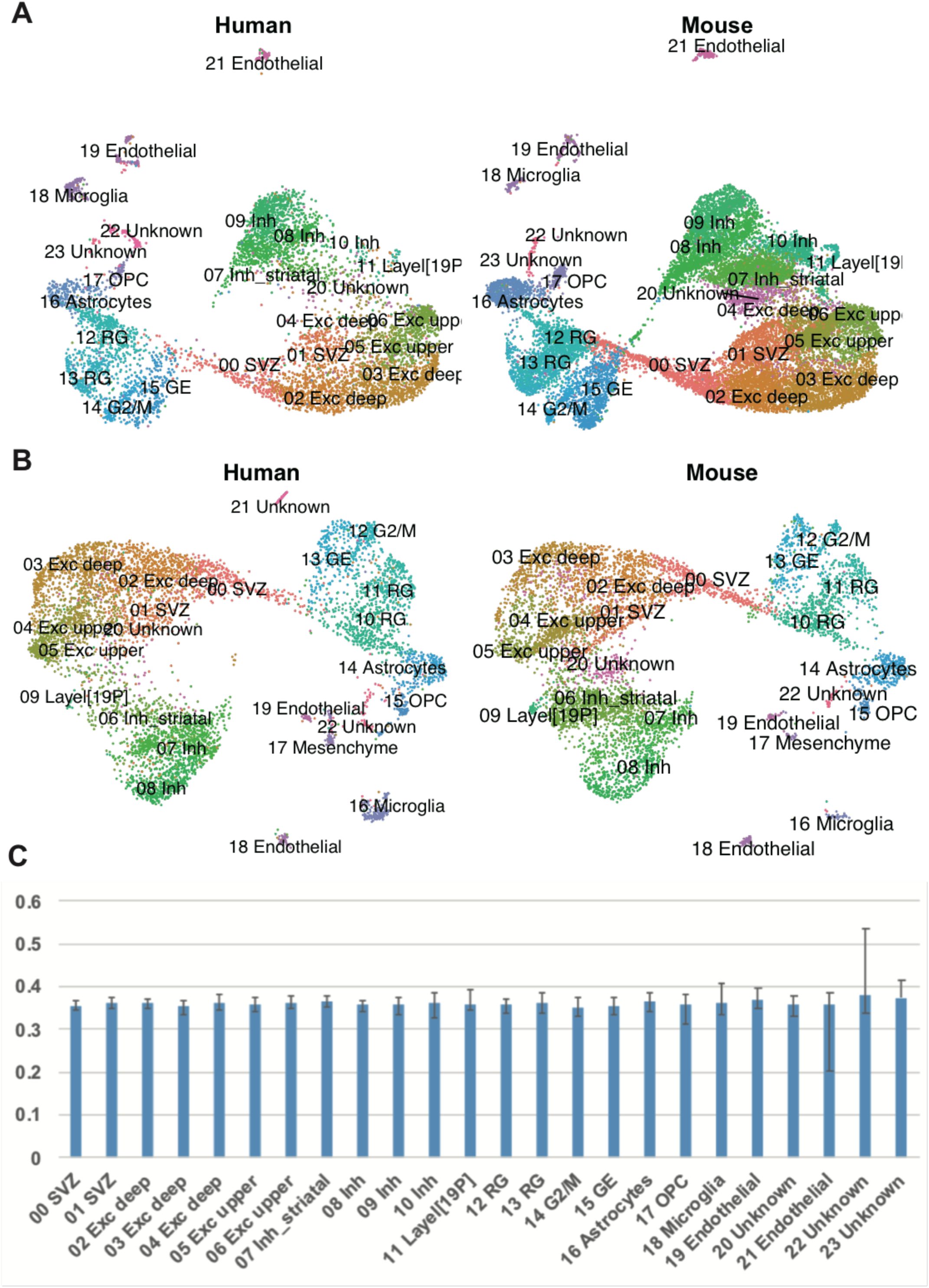

**Figure S2.**
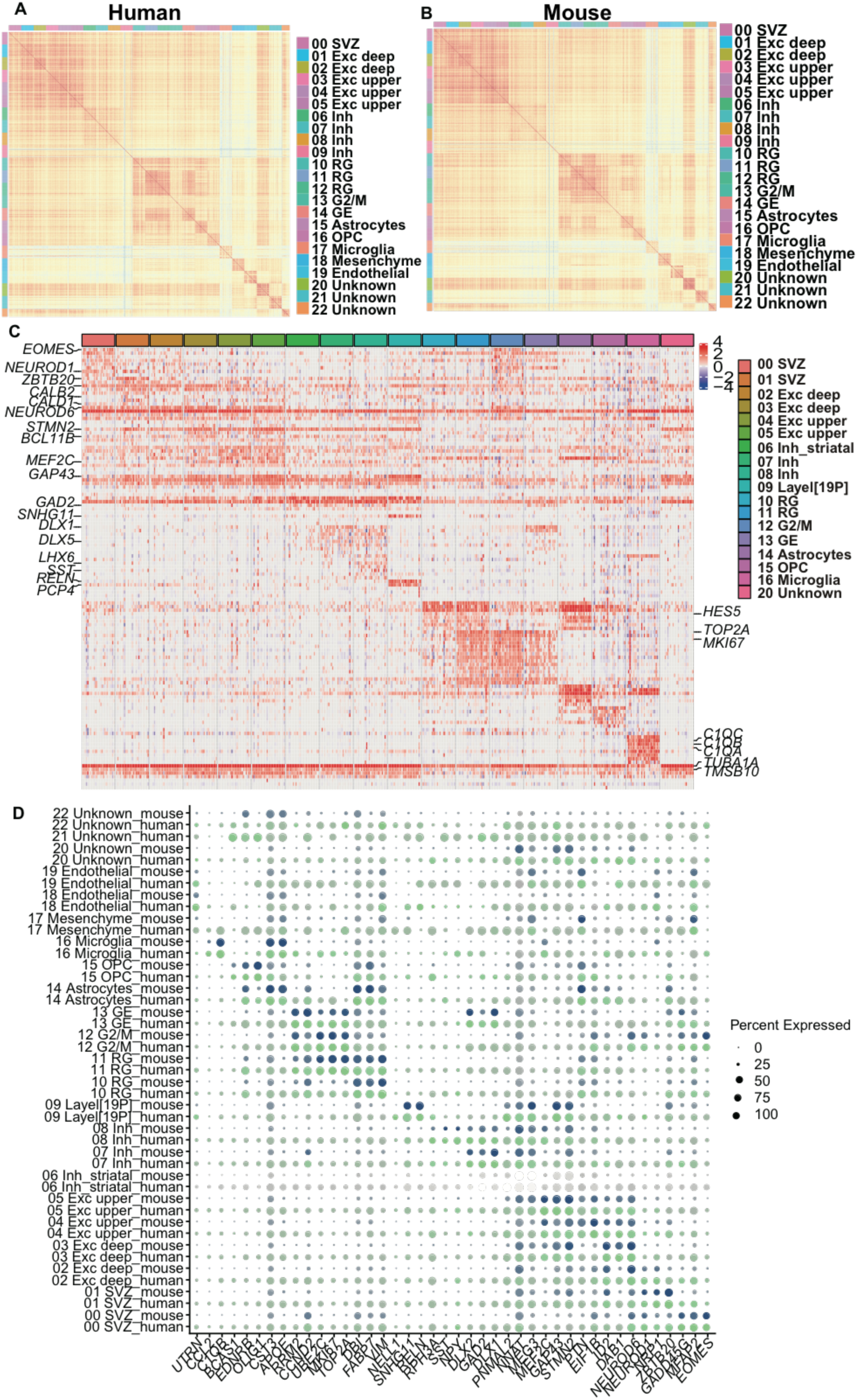

**Figure S3.**
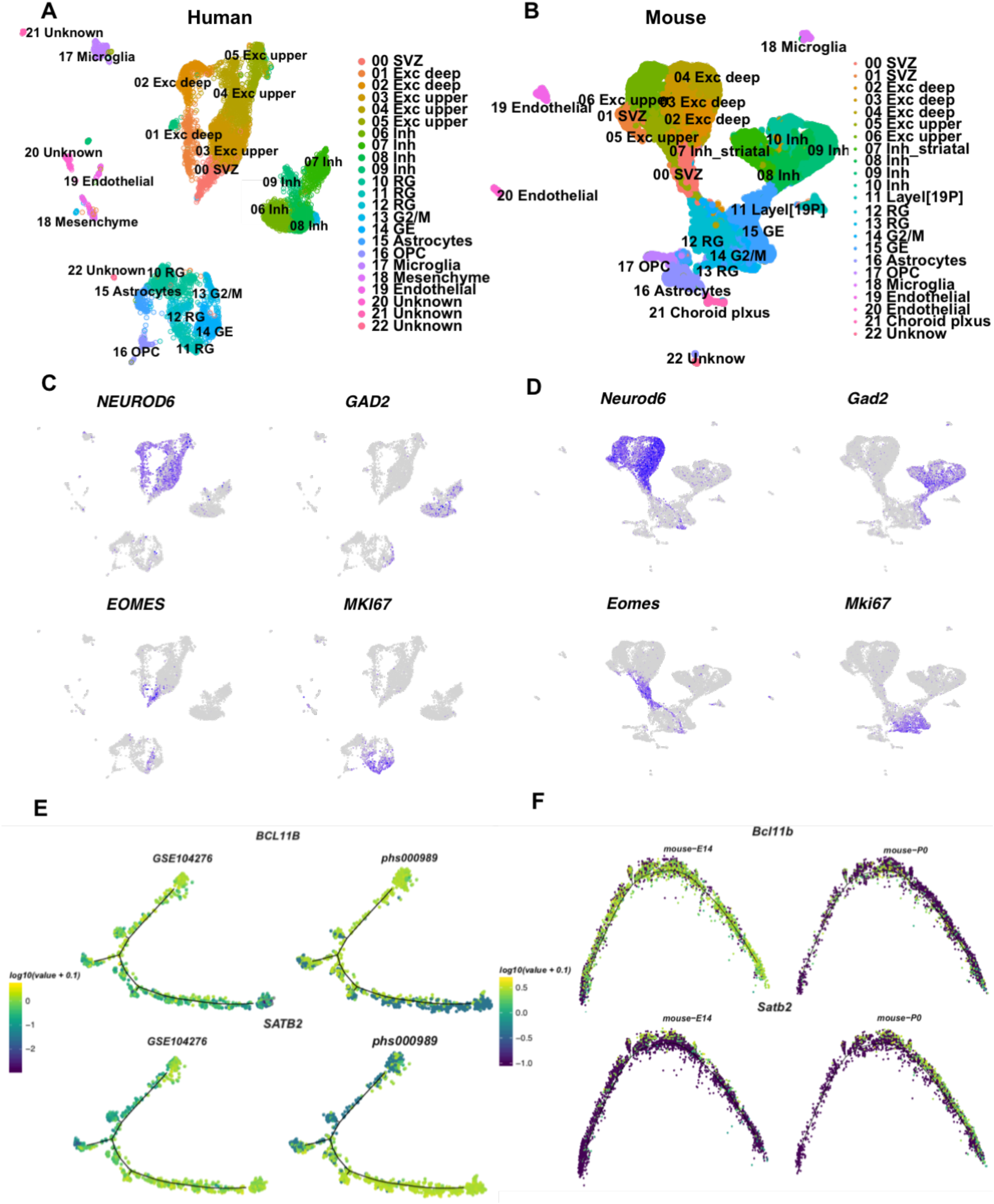
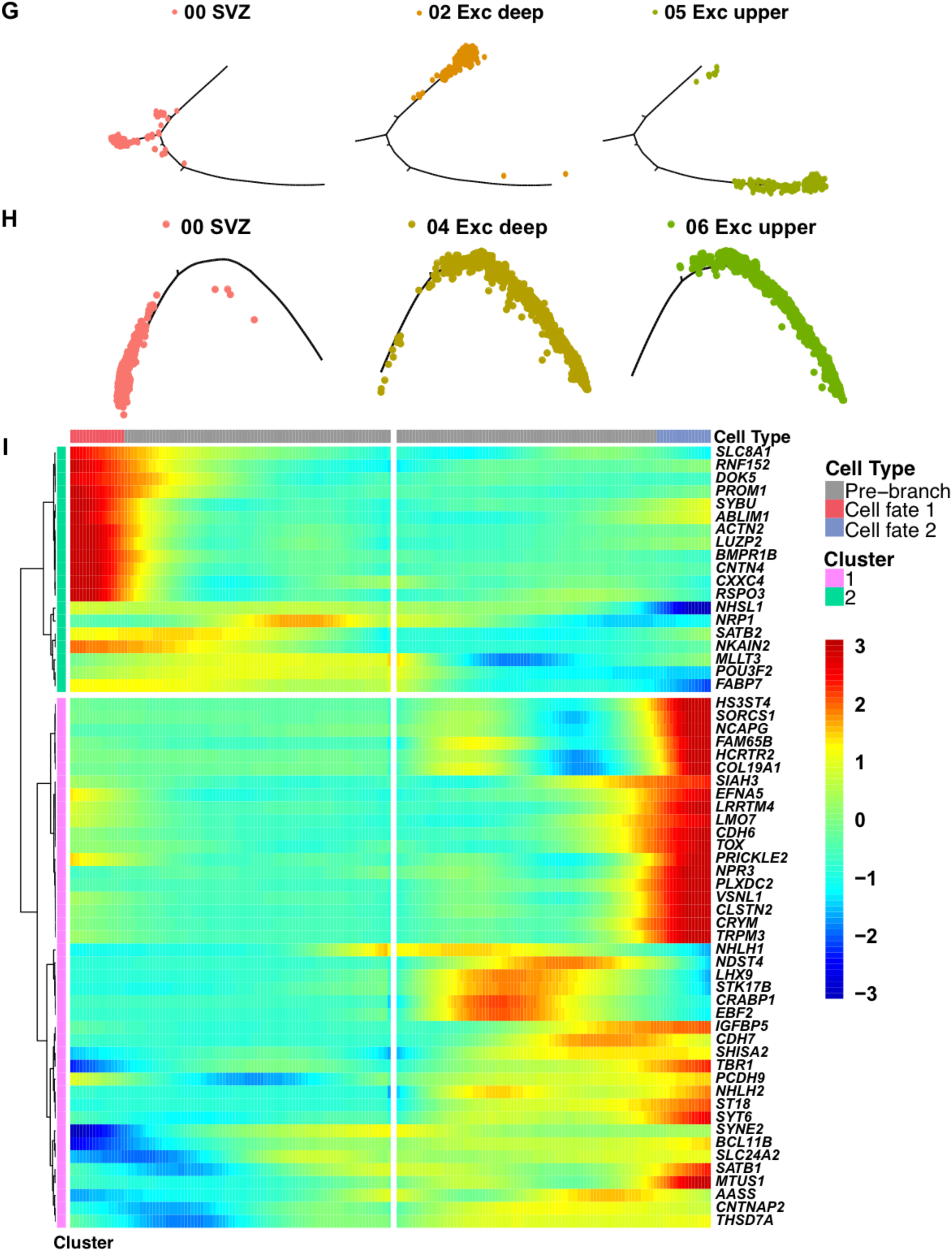
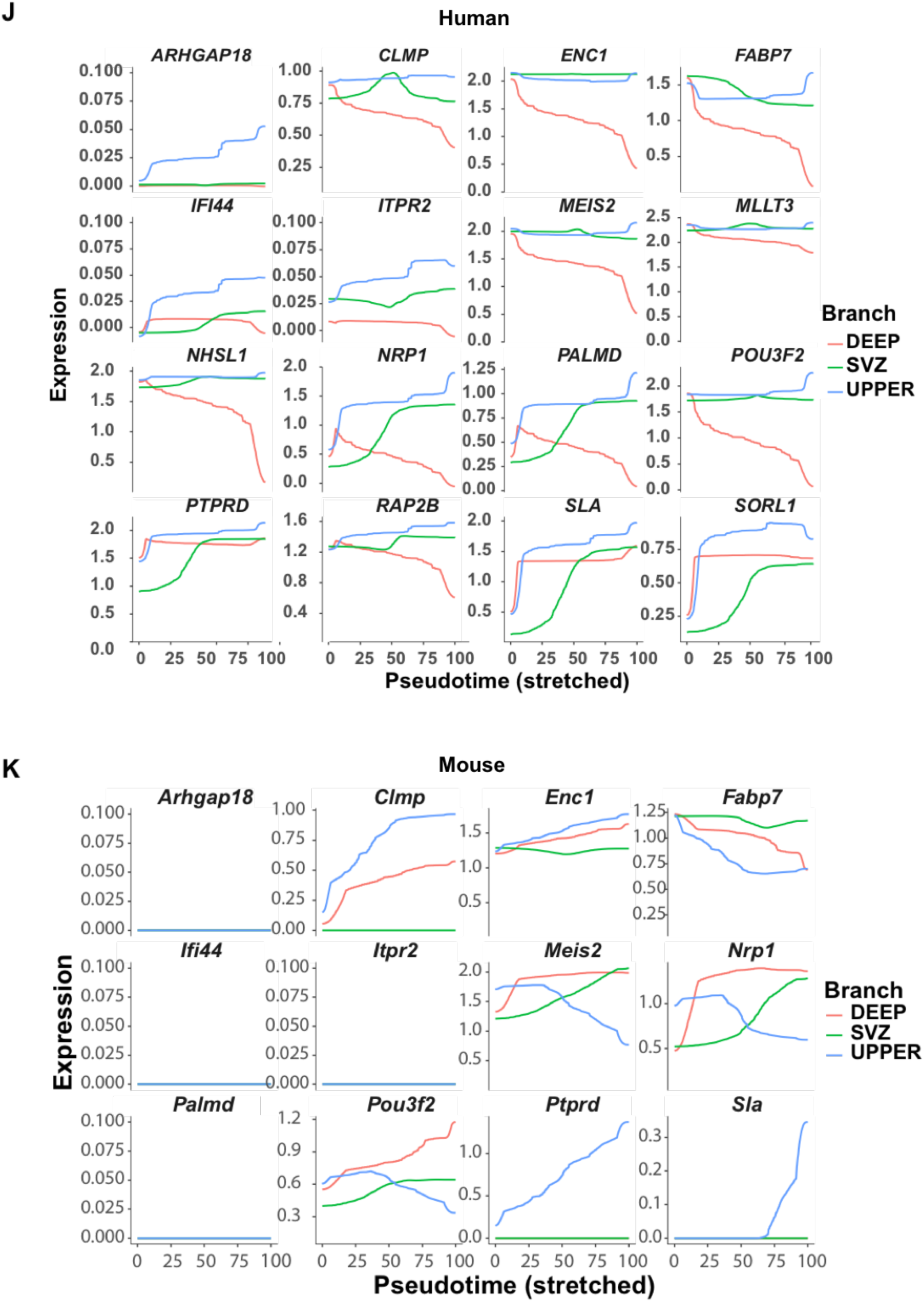

**Figure S4.**
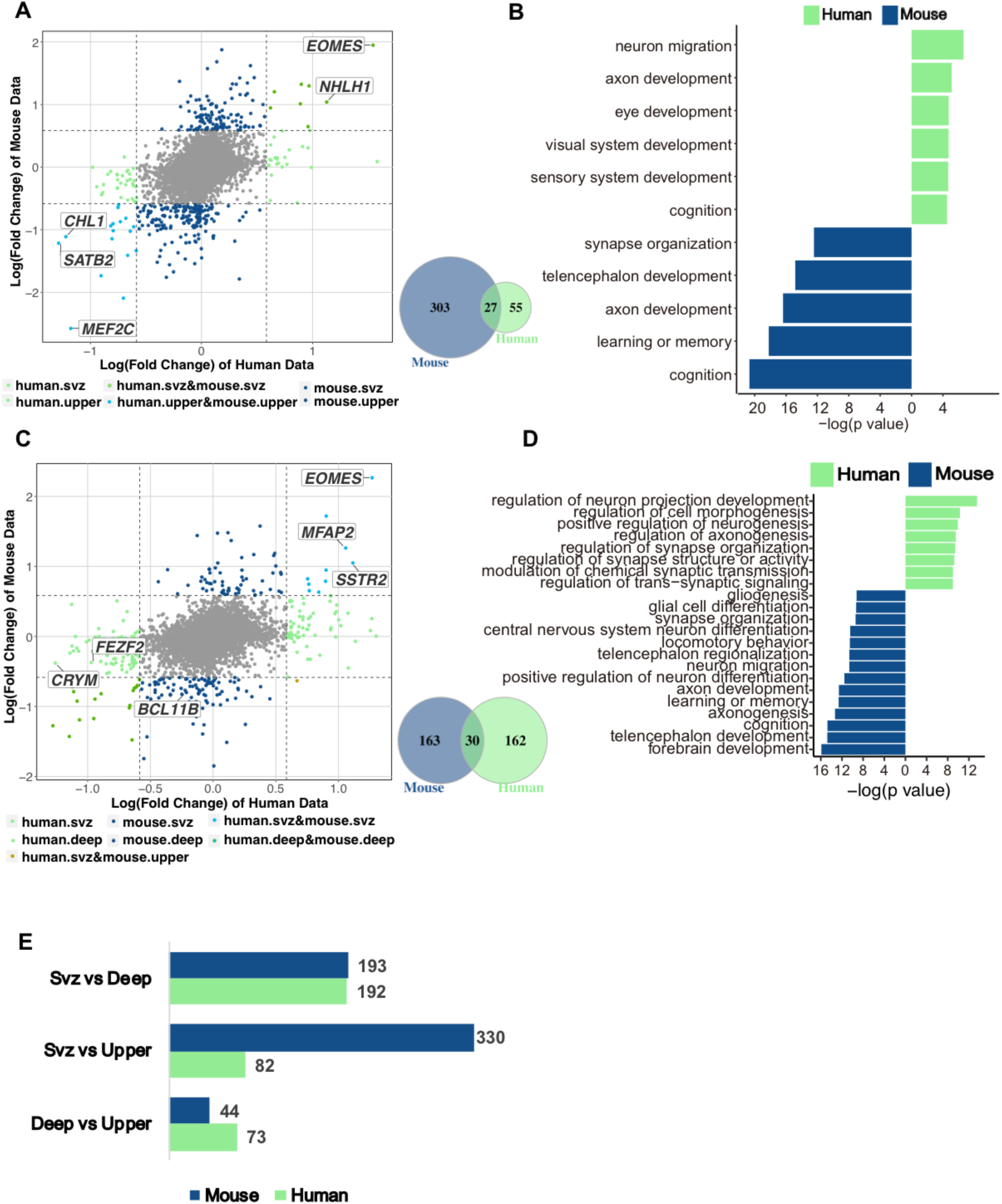

**Figure S5.**
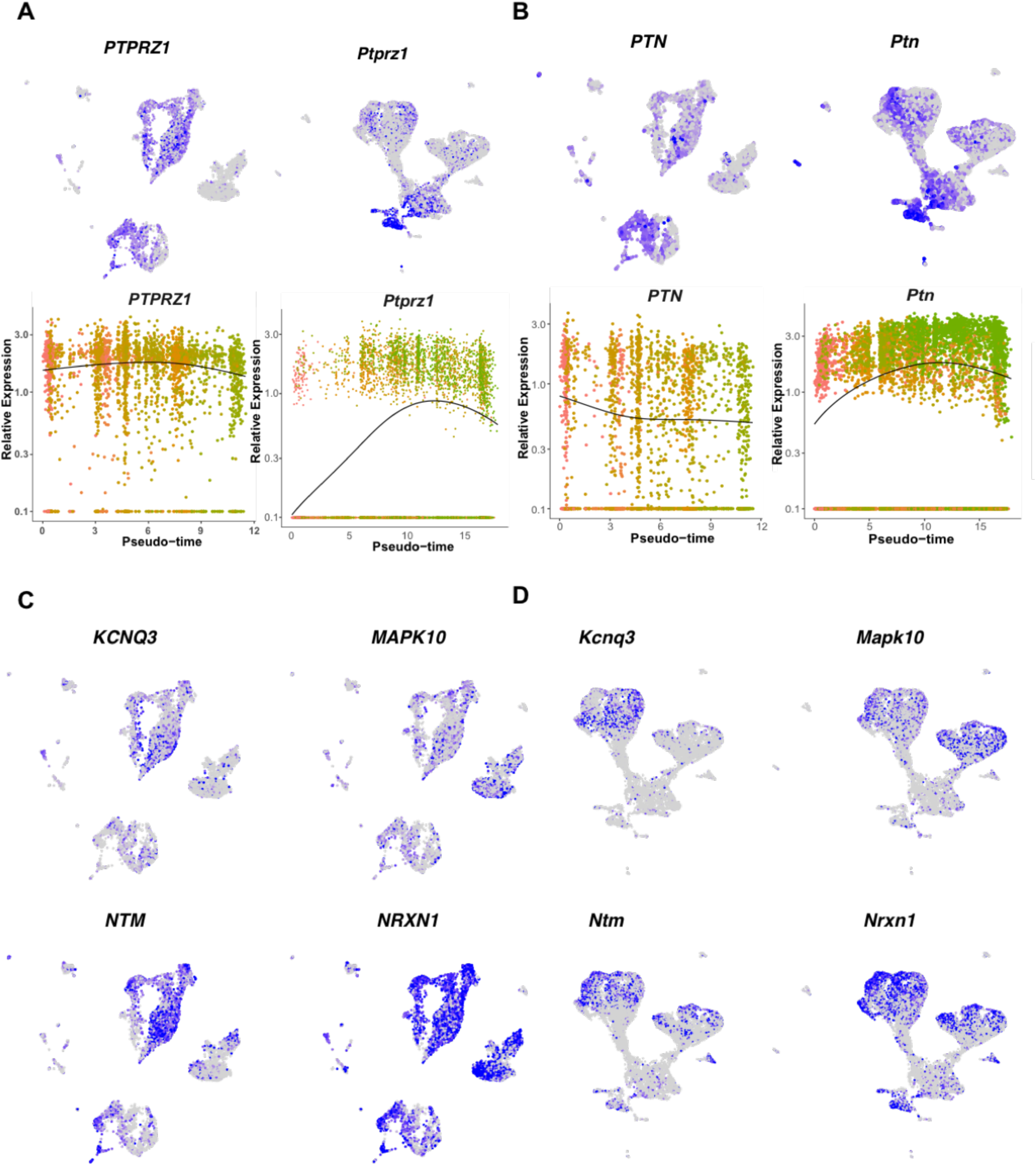
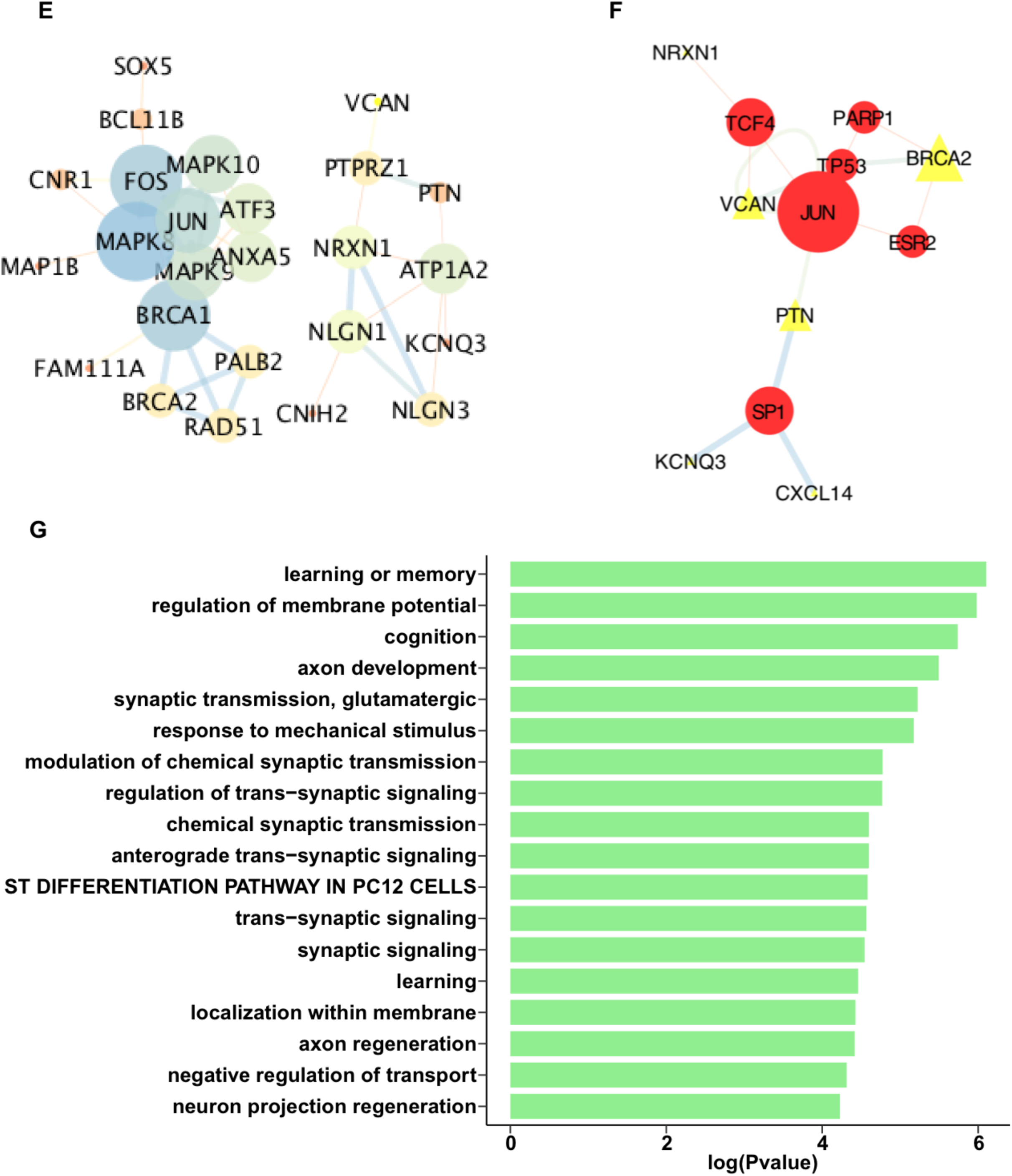

**Figure S6.**
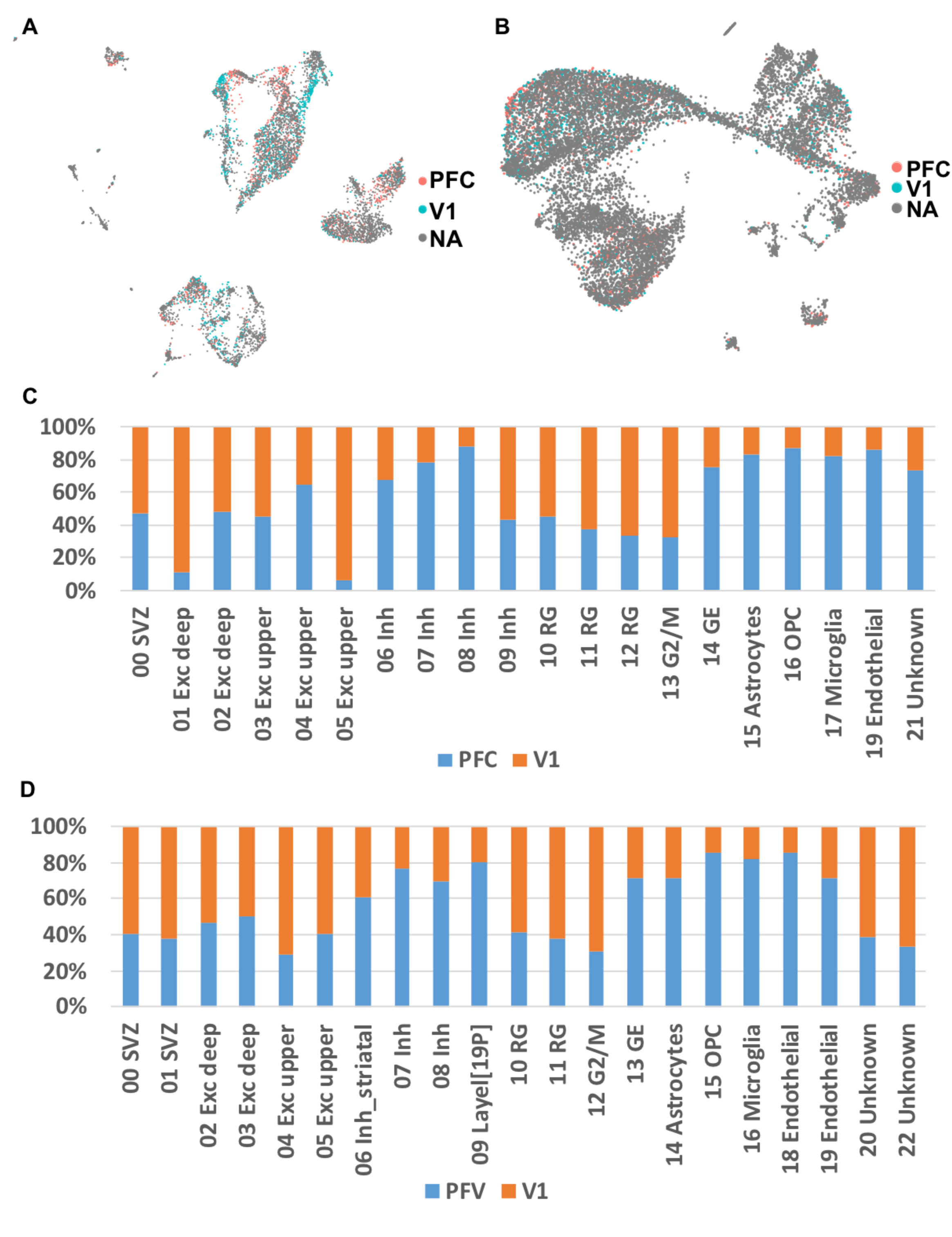

**Figure S7.**
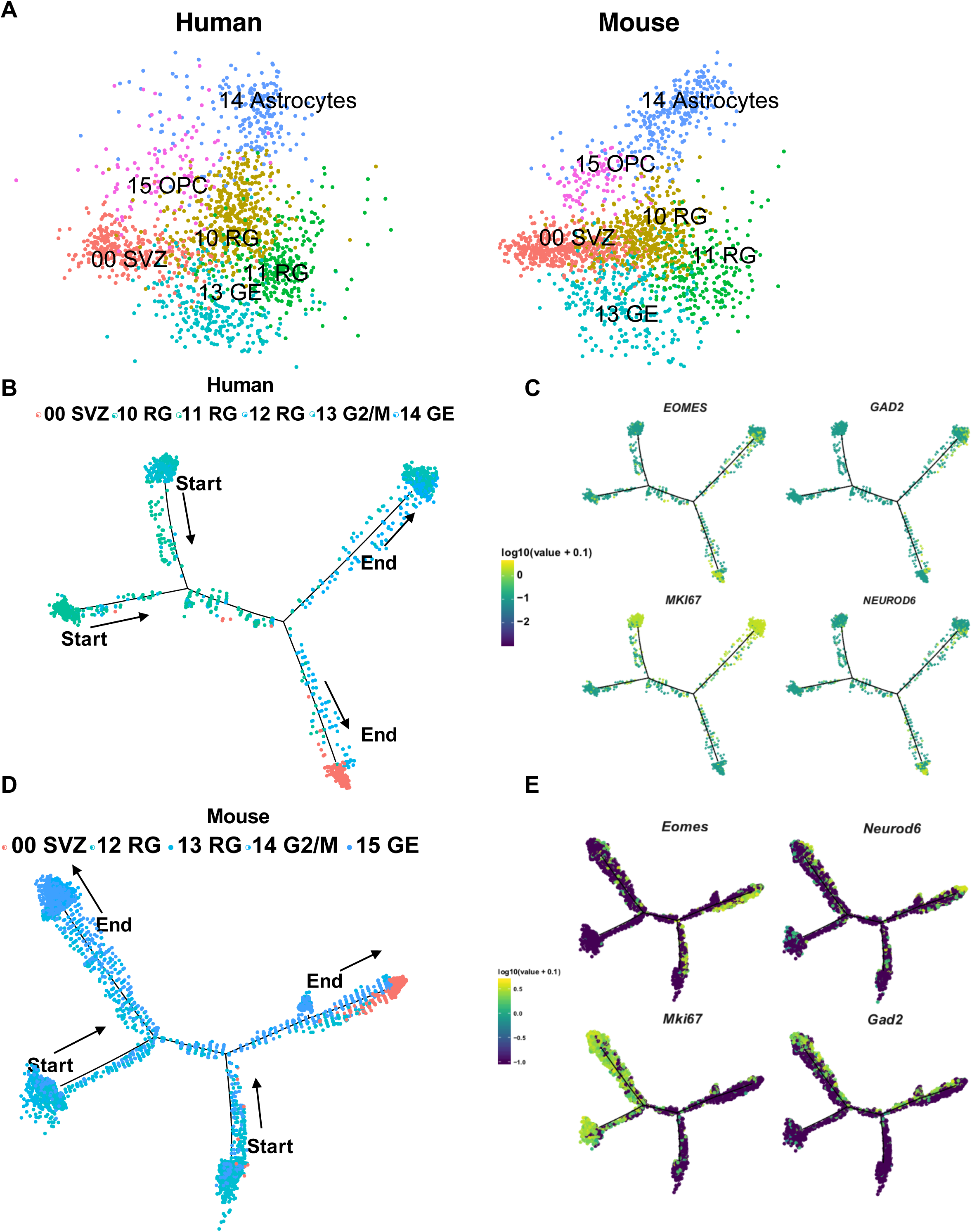
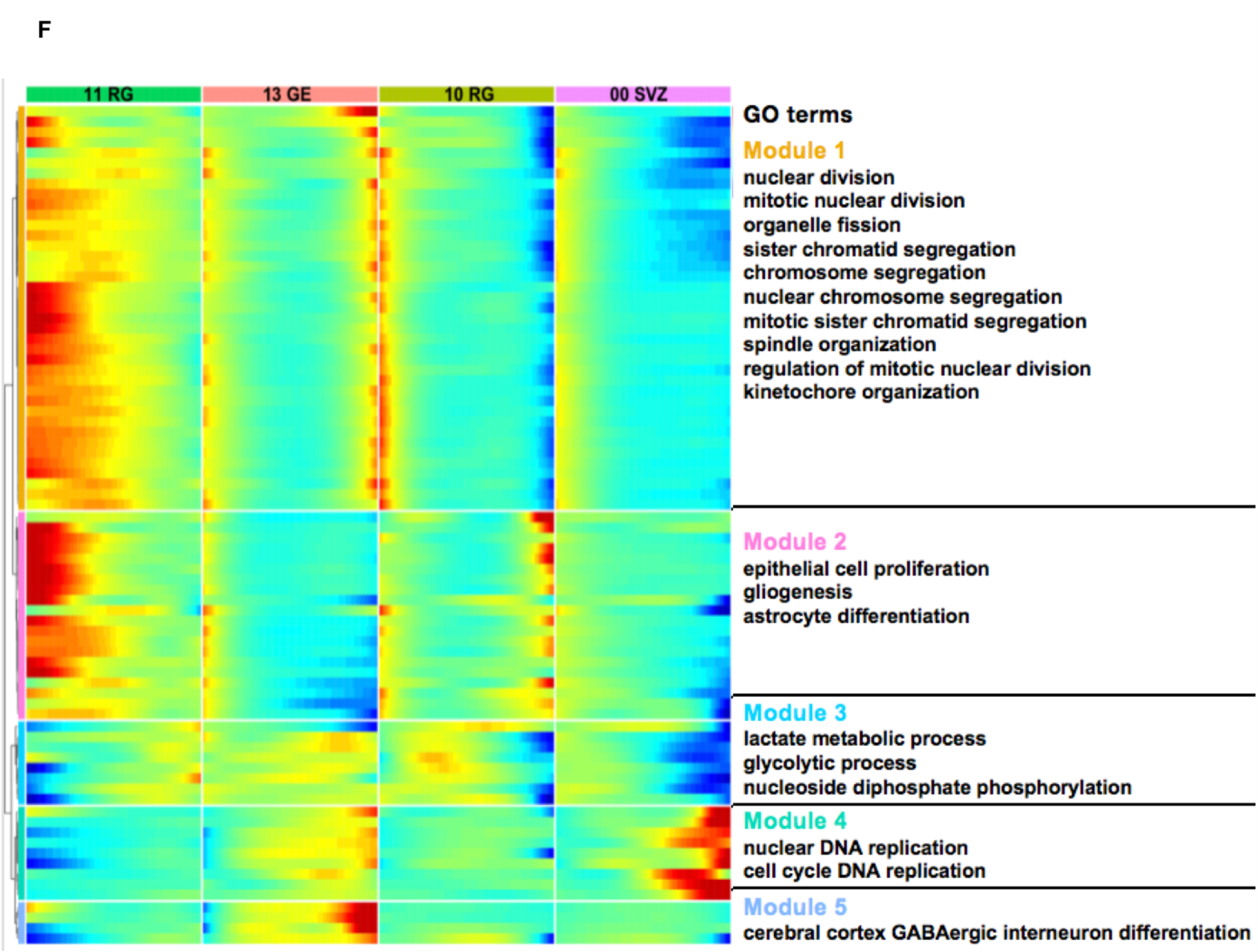

**Figure S8.**
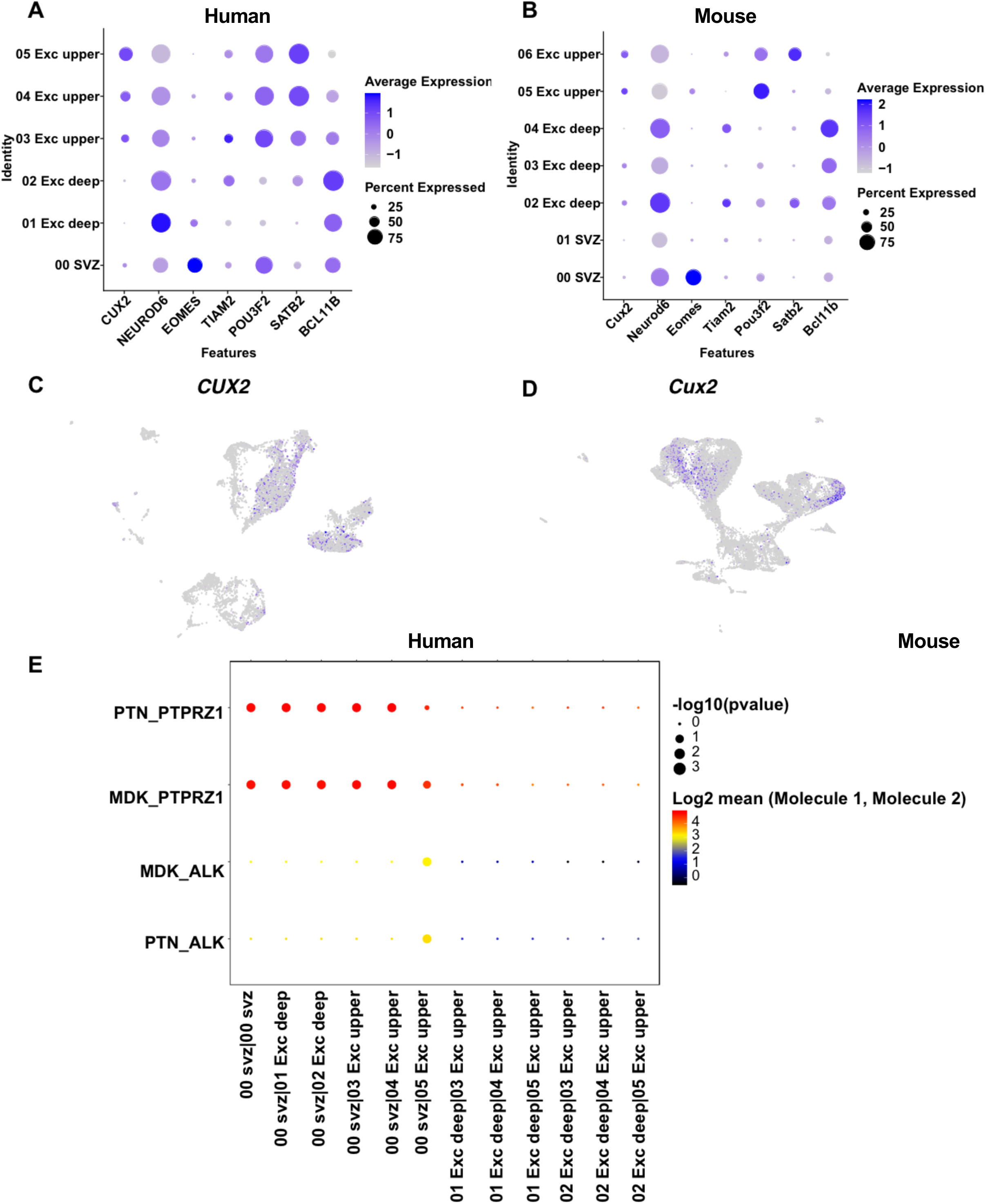
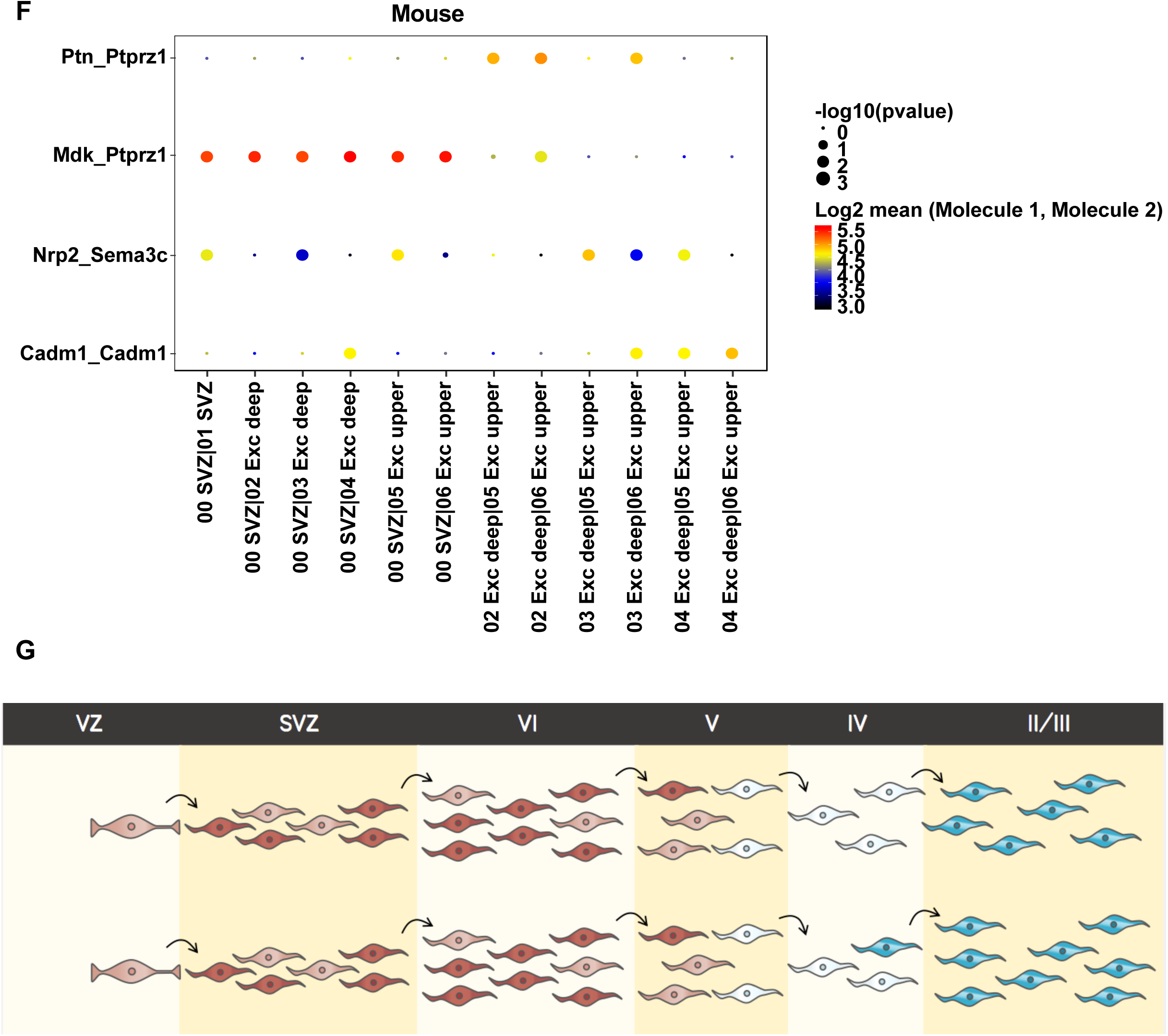

## Notes

### Competing Interest Statement

The authors have declared no competing interest.

## REFERENCES

1. Otis EM, Brent R: Equivalent ages in mouse and human embryos. The Anatomical record 1954, 120:33–63.

2. Workman AD, Charvet CJ, Clancy B, Darlington RB, Finlay BL: Modeling transformations of neurodevelopmental sequences across mammalian species. Journal of Neuroscience 2013, 33:7368–7383.

3. Wichterle H, Turnbull DH, Nery S, Fishell G, Alvarez-Buylla A: In utero fate mapping reveals distinct migratory pathways and fates of neurons born in the mammalian basal forebrain. Development 2001, 128:3759–3771.

4. Cobos I, Puelles L, Martínez S: The avian telencephalic subpallium originates inhibitory neurons that invade tangentially the pallium (dorsal ventricular ridge and cortical areas). Developmental biology 2001, 239:30–45.

5. Wonders CP, Anderson SA: The origin and specification of cortical interneurons. Nature Reviews Neuroscience 2006, 7:687–696.

6. Guo J, Anton E: Decision making during interneuron migration in the developing cerebral cortex. Trends in cell biology 2014, 24:342–351.

7. Gorski JA, Talley T, Qiu M, Puelles L, Rubenstein JL, Jones KR: Cortical excitatory neurons and glia, but not GABAergic neurons, are produced in the Emx1-expressing lineage. Journal of Neuroscience 2002, 22:6309–6314.

8. Molyneaux BJ, Arlotta P, Menezes JR, Macklis JD: Neuronal subtype specification in the cerebral cortex. Nature reviews neuroscience 2007, 8:427–437.

9. Nowakowski TJ, Bhaduri A, Pollen AA, Alvarado B, Mostajo-Radji MA, Di Lullo E, Haeussler M, Sandoval-Espinosa C, Liu SJ, Velmeshev D: Spatiotemporal gene expression trajectories reveal developmental hierarchies of the human cortex. Science 2017, 358:1318–1323.

10. Hodge RD, Bakken TE, Miller JA, Smith KA, Barkan ER, Graybuck LT, Close JL, Long B, Johansen N, Penn O: Conserved cell types with divergent features in human versus mouse cortex. Nature 2019, 573:61–68.

11. Molnár Z, Métin C, Stoykova A, Tarabykin V, Price DJ, Francis F, Meyer G, Dehay C, Kennedy H: Comparative aspects of cerebral cortical development. European Journal of Neuroscience 2006, 23:921–934.

12. Monuki ES, Walsh CA: Mechanisms of cerebral cortical patterning in mice and humans. Nature neuroscience 2001, 4:1199–1206.

13. Yuan G-C, Cai L, Elowitz M, Enver T, Fan G, Guo G, Irizarry R, Kharchenko P, Kim J, Orkin S: Challenges and emerging directions in single-cell analysis. Genome biology 2017, 18:84.

14. Poirion OB, Zhu X, Ching T, Garmire L: Single-cell transcriptomics bioinformatics and computational challenges. Frontiers in genetics 2016, 7:163.

15. Papalexi E, Satija R: Single-cell RNA sequencing to explore immune cell heterogeneity. Nature Reviews Immunology 2018, 18:35.

16. Giladi A, Amit I: Single-cell genomics: a stepping stone for future immunology discoveries. Cell 2018, 172:14–21.

17. Zhong S, Zhang S, Fan X, Wu Q, Yan L, Dong J, Zhang H, Li L, Sun L, Pan N: A single-cell RNA-seq survey of the developmental landscape of the human prefrontal cortex. Nature 2018, 555:524–528.

18. Loo L, Simon JM, Xing L, McCoy ES, Niehaus JK, Guo J, Anton E, Zylka MJ: Single-cell transcriptomic analysis of mouse neocortical development. Nature communications 2019, 10:1–11.

19. McInnes L, Healy J, Melville J: Umap: Uniform manifold approximation and projection for dimension reduction. arXiv preprint 180203426 2018.

20. Camp JG, Badsha F, Florio M, Kanton S, Gerber T, Wilsch-Bräuninger M, Lewitus E, Sykes A, Hevers W, Lancaster M: Human cerebral organoids recapitulate gene expression programs of fetal neocortex development. Proceedings of the National Academy of Sciences 2015, 112:15672–15677.

21. Han X, Wang R, Zhou Y, Fei L, Sun H, Lai S, Saadatpour A, Zhou Z, Chen H, Ye F: Mapping the mouse cell atlas by microwell-seq. Cell 2018, 172:1091-1107. e1017.

22. Englund C, Fink A, Lau C, Pham D, Daza RA, Bulfone A, Kowalczyk T, Hevner RF: Pax6, Tbr2, and Tbr1 are expressed sequentially by radial glia, intermediate progenitor cells, and postmitotic neurons in developing neocortex. Journal of Neuroscience 2005, 25:247–251.

23. Nagao M, Ogata T, Sawada Y, Gotoh Y: Zbtb20 promotes astrocytogenesis during neocortical development. Nature communications 2016, 7:1–14.

24. Nielsen JV, Thomassen M, Møllgård K, Noraberg J, Jensen NA: Zbtb20 defines a hippocampal neuronal identity through direct repression of genes that control projection neuron development in the isocortex. Cerebral Cortex 2014, 24:1216–1229.

25. Doeppner TR, Herz J, Bähr M, Tonchev AB, Stoykova A: Zbtb20 regulates developmental neurogenesis in the olfactory bulb and gliogenesis after adult brain injury. Molecular neurobiology 2019, 56:567–582.

26. Kohwi M, Petryniak MA, Long JE, Ekker M, Obata K, Yanagawa Y, Rubenstein JL, Alvarez-Buylla A: A subpopulation of olfactory bulb GABAergic interneurons is derived from Emx1-and Dlx5/6-expressing progenitors. Journal of Neuroscience 2007, 27:6878–6891.

27. Arnold SJ, Huang G-J, Cheung AF, Era T, Nishikawa S-I, Bikoff EK, Molnár Z, Robertson EJ, Groszer M: The T-box transcription factor Eomes/Tbr2 regulates neurogenesis in the cortical subventricular zone. Genes & development 2008, 22:2479–2484.

28. Greig LC, Woodworth MB, Galazo MJ, Padmanabhan H, Macklis JD: Molecular logic of neocortical projection neuron specification, development and diversity. Nature Reviews Neuroscience 2013, 14:755–769.

29. Leifer D, Krainc D, Yu Y-T, McDermott J, Breitbart RE, Heng J, Neve RL, Kosofsky B, Nadal-Ginard B, Lipton SA: MEF2C, a MADS/MEF2-family transcription factor expressed in a laminar distribution in cerebral cortex. Proceedings of the National Academy of Sciences 1993, 90:1546–1550.

30. Tomita K, Gotoh H, Tomita K, Yamauchi N, Sanbo M: Multiple patterns of spatiotemporal changes in layer-specific gene expression in the developing visual cortex of higher mammals. Neuroscience research 2012, 73:207–217.

31. Mayer C, Hafemeister C, Bandler RC, Machold R, Brito RB, Jaglin X, Allaway K, Butler A, Fishell G, Satija R: Developmental diversification of cortical inhibitory interneurons. Nature 2018, 555:457–462.

32. Tasic B, Menon V, Nguyen TN, Kim TK, Jarsky T, Yao Z, Levi B, Gray LT, Sorensen SA, Dolbeare T: Adult mouse cortical cell taxonomy revealed by single cell transcriptomics. Nature neuroscience 2016, 19:335.

33. Mi D, Li Z, Lim L, Li M, Moissidis M, Yang Y, Gao T, Hu TX, Pratt T, Price DJ: Early emergence of cortical interneuron diversity in the mouse embryo. Science 2018, 360:81–85.

34. Ghanem N, Yu M, Long J, Hatch G, Rubenstein JL, Ekker M: Distinct cis-regulatory elements from the Dlx1/Dlx2 locus mark different progenitor cell populations in the ganglionic eminences and different subtypes of adult cortical interneurons. Journal of Neuroscience 2007, 27:5012–5022.

35. Donovan SL, Mamounas LA, Andrews AM, Blue ME, McCasland JS: GAP-43 is critical for normal development of the serotonergic innervation in forebrain. Journal of Neuroscience 2002, 22:3543–3552.

36. Desplats PA, Lambert JR, Thomas EA: Functional roles for the striatal-enriched transcription factor, Bcl11b, in the control of striatal gene expression and transcriptional dysregulation in Huntington’s disease. Neurobiology of disease 2008, 31:298–308.

37. Geirsdottir L, David E, Keren-Shaul H, Weiner A, Bohlen SC, Neuber J, Balic A, Giladi A, Sheban F, Dutertre C-A: Cross-Species Single-Cell Analysis Reveals Divergence of the Primate Microglia Program. Cell 2019, 179:1609-1622. e1616.

38. Jansen IE, Savage JE, Watanabe K, Bryois J, Williams DM, Steinberg S, Sealock J, Karlsson IK, Hägg S, Athanasiu L: Genome-wide meta-analysis identifies new loci and functional pathways influencing Alzheimer’s disease risk. Nature genetics 2019, 51:404–413.

39. Lambert J, Ibrahim-Verbaas C, Harold D, Naj A, Sims R, Bellenguez C, DeStafano A, Bis J, Beecham G, Grenier-Boley B: European Alzheimer’s Disease Initiative (EADI); Genetic and Environmental Risk in Alzheimer’s Disease; Alzheimer’s Disease Genetic Consortium; Cohorts for Heart and Aging Research in Genomic Epidemiology. Meta-analysis of 74,046 individuals identifies 11 new susceptibility loci for Alzheimer’s disease. Nat Genet 2013, 45:1452–1458.

40. Keren-Shaul H, Spinrad A, Weiner A, Matcovitch-Natan O, Dvir-Szternfeld R, Ulland TK, David E, Baruch K, Lara-Astaiso D, Toth B: A unique microglia type associated with restricting development of Alzheimer’s disease. Cell 2017, 169:1276-1290. e1217.

41. Lim DA, Alvarez-Buylla A: The adult ventricular–subventricular zone (V-SVZ) and olfactory bulb (OB) neurogenesis. Cold Spring Harbor perspectives in biology 2016, 8:a018820.

42. Wang T-W, Zhang H, Parent JM: Retinoic acid regulates postnatal neurogenesis in the murine subventricular zone-olfactory bulb pathway. Development 2005, 132:2721–2732.

43. O’rourke NA, Weiler NC, Micheva KD, Smith SJ: Deep molecular diversity of mammalian synapses: why it matters and how to measure it. Nature Reviews Neuroscience 2012, 13:365–379.

44. Fantin A, Maden CH, Ruhrberg C: Neuropilin ligands in vascular and neuronal patterning. Portland Press Ltd.; 2009.

45. Lipina TV, Prasad T, Yokomaku D, Luo L, Connor SA, Kawabe H, Wang YT, Brose N, Roder JC, Craig AM: Cognitive deficits in calsyntenin-2-deficient mice associated with reduced GABAergic transmission. Neuropsychopharmacology 2016, 41:802.

46. Efremova M, Vento-Tormo M, Teichmann SA, Vento-Tormo R: CellPhoneDB v2. 0: Inferring cell-cell communication from combined expression of multi-subunit receptor-ligand complexes. bioRxiv 2019: 680926.

47. Janoueix-Lerosey I, Lopez-Delisle L, Delattre O, Rohrer H: The ALK receptor in sympathetic neuron development and neuroblastoma. Cell and tissue research 2018, 372:325–337.

48. Motegi A, Fujimoto J, Kotani M, Sakuraba H, Yamamoto T: ALK receptor tyrosine kinase promotes cell growth and neurite outgrowth. Journal of cell science 2004, 117:3319–3329.

49. Wellstein A: ALK receptor activation, ligands and therapeutic targeting in glioblastoma and in other cancers. Frontiers in oncology 2012, 2:192.

50. Herradon G, Pérez - Garcí a C: Targeting midkine and pleiotrophin signalling pathways in addiction and neurodegenerative disorders: recent progress and perspectives. British journal of pharmacology 2014, 171:837–848.

51. Rusanescu G, Mao J: Notch3 is necessary for neuronal differentiation and maturation in the adult spinal cord. Journal of cellular and molecular medicine 2014, 18:2103–2116.

52. Dvoriantchikova G, Perea-Martinez I, Pappas S, Barry AF, Danek D, Dvoriantchikova X, Pelaez D, Ivanov D: Molecular characterization of notch1 positive progenitor cells in the developing retina. PloS one 2015, 10.

53. Palmer A, Klein R: Multiple roles of ephrins in morphogenesis, neuronal networking, and brain function. Genes & development 2003, 17:1429–1450.

54. Yuferov V, Ho A, Morgello S, Yang Y, Ott J, Kreek MJ: Expression of ephrin receptors and ligands in postmortem brains of HIV-infected subjects with and without cognitive impairment. Journal of Neuroimmune Pharmacology 2013, 8:333–344.

55. Paganoni S, Bernstein J, Ferreira A: Ror1-Ror2 complexes modulate synapse formation in hippocampal neurons. Neuroscience 2010, 165:1261–1274.

56. Hansell CA, MacLellan LM, Oldham RS, Doonan J, Chapple KJ, Anderson EJ, Linington C, McInnes IB, Nibbs RJ, Goodyear CS: The atypical chemokine receptor ACKR2 suppresses Th17 responses to protein autoantigens. Immunology and cell biology 2015, 93:167–176.

57. Harkany T, Keimpema E, Barabás K, Mulder J: Endocannabinoid functions controlling neuronal specification during brain development. Molecular and cellular endocrinology 2008, 286:S84–S90.

58. Mulder J, Aguado T, Keimpema E, Barabás K, Rosado CJB, Nguyen L, Monory K, Marsicano G, Di Marzo V, Hurd YL: Endocannabinoid signaling controls pyramidal cell specification and long-range axon patterning. Proceedings of the National Academy of Sciences 2008, 105:8760–8765.

59. Watson S, Chambers D, Hobbs C, Doherty P, Graham A: The endocannabinoid receptor, CB1, is required for normal axonal growth and fasciculation. Molecular and Cellular Neuroscience 2008, 38:89–97.

60. Wu CS, Zhu J, Wager-Miller J, Wang S, O’Leary D, Monory K, Lutz B, Mackie K, Lu HC: Requirement of cannabinoid CB1 receptors in cortical pyramidal neurons for appropriate development of corticothalamic and thalamocortical projections. European Journal of Neuroscience 2010, 32:693–706.

61. Keimpema E, Barabas K, Morozov YM, Tortoriello G, Torii M, Cameron G, Yanagawa Y, Watanabe M, Mackie K, Harkany T: Differential subcellular recruitment of monoacylglycerol lipase generates spatial specificity of 2-arachidonoyl glycerol signaling during axonal pathfinding. Journal of Neuroscience 2010, 30:13992–14007.

62. Mackie K: Signaling via CNS cannabinoid receptors. Molecular and cellular endocrinology 2008, 286:S60–S65.

63. Bilkei-Gorzo A, Racz I, Valverde O, Otto M, Michel K, Sarstre M, Zimmer A: Early age-related cognitive impairment in mice lacking cannabinoid CB1 receptors. Proceedings of the National Academy of Sciences 2005, 102:15670–15675.

64. Schneider M, Kasanetz F, Lynch DL, Friemel CM, Lassalle O, Hurst DP, Steindel F, Monory K, Schäfer C, Miederer I: Enhanced functional activity of the cannabinoid type-1 receptor mediates adolescent behavior. Journal of Neuroscience 2015, 35:13975–13988.

65. Juhász G, Csepany E, Magyar M, Édes AE, Eszlári N, Hullam G, Antal P, Kokonyei G, Anderson I, Deakin J: Variants in the CNR1 gene predispose to headache with nausea in the presence of life stress. Genes, Brain and Behavior 2017, 16:384–393.

66. Jandrig B, Seitz S, Hinzmann B, Arnold W, Micheel B, Koelble K, Siebert R, Schwartz A, Ruecker K, Schlag PM: ST18 is a breast cancer tumor suppressor gene at human chromosome 8q11. 2. Oncogene 2004, 23:9295–9302.

67. Raivich G, Bohatschek M, Da Costa C, Iwata O, Galiano M, Hristova M, Nateri AS, Makwana M, Riera-Sans Ls, Wolfer DP: The AP-1 transcription factor c-Jun is required for efficient axonal regeneration. Neuron 2004, 43:57–67.

68. Yamasaki T, Kawasaki H, Nishina H: Diverse roles of JNK and MKK pathways in the brain. Journal of signal transduction 2012, 2012.

69. Tang C, Wang M, Wang P, Wang L, Wu Q, Guo W: Neural stem cells behave as a functional niche for the maturation of newborn neurons through the secretion of PTN. Neuron 2019, 101:32-44. e36.

70. González-Castillo C, Ortuño-Sahagún D, Guzmán-Brambila C, Pallàs M, Rojas-Mayorquín AE: Pleiotrophin as a central nervous system neuromodulator, evidences from the hippocampus. Frontiers in cellular neuroscience 2015, 8:443.

71. Lu KV, Jong KA, Kim GY, Singh J, Dia EQ, Yoshimoto K, Wang MY, Cloughesy TF, Nelson SF, Mischel PS: Differential induction of glioblastoma migration and growth by two forms of pleiotrophin. Journal of Biological Chemistry 2005, 280:26953–26964.

72. Szklarczyk D, Gable AL, Lyon D, Junge A, Wyder S, Huerta-Cepas J, Simonovic M, Doncheva NT, Morris JH, Bork P: STRING v11: protein–protein association networks with increased coverage, supporting functional discovery in genome-wide experimental datasets. Nucleic acids research 2019, 47:D607–D613.

73. Castiglioni V, Faedo A, Onorati M, Bocchi VD, Li Z, Iennaco R, Vuono R, Bulfamante GP, Muzio L, Martino G: Dynamic and cell-specific DACH1 expression in human neocortical and striatal development. Cerebral Cortex 2019, 29:2115–2124.

74. Zimmer C, Tiveron M-C, Bodmer R, Cremer H: Dynamics of Cux2 expression suggests that an early pool of SVZ precursors is fated to become upper cortical layer neurons. Cerebral cortex 2004, 14:1408–1420.

75. Cubelos B, Sebastián-Serrano A, Beccari L, Calcagnotto ME, Cisneros E, Kim S, Dopazo A, Alvarez-Dolado M, Redondo JM, Bovolenta P: Cux1 and Cux2 regulate dendritic branching, spine morphology, and synapses of the upper layer neurons of the cortex. Neuron 2010, 66:523–535.

76. Krellman JW, Ruiz HH, Marciano VA, Mondrow B, Croll SD: Behavioral and neuroanatomical abnormalities in pleiotrophin knockout mice. PloS one 2014, 9.

77. Stuart T, Butler A, Hoffman P, Hafemeister C, Papalexi E, Mauck III WM, Hao Y, Stoeckius M, Smibert P, Satija R: Comprehensive integration of single-cell data. Cell 2019, 177:1888-1902. e1821.

78. Trapnell C, Cacchiarelli D, Grimsby J, Pokharel P, Li S, Morse M, Lennon NJ, Livak KJ, Mikkelsen TS, Rinn JL: The dynamics and regulators of cell fate decisions are revealed by pseudotemporal ordering of single cells. Nature biotechnology 2014, 32:381.

79. Yu G, Wang L-G, Han Y, He Q-Y: clusterProfiler: an R package for comparing biological themes among gene clusters. Omics: a journal of integrative biology 2012, 16:284–287.

80. Zhou Y, Zhou B, Pache L, Chang M, Khodabakhshi AH, Tanaseichuk O, Benner C, Chanda SK: Metascape provides a biologist-oriented resource for the analysis of systems-level datasets. Nature communications 2019, 10:1–10.

81. Shannon P, Markiel A, Ozier O, Baliga NS, Wang JT, Ramage D, Amin N, Schwikowski B, Ideker T: Cytoscape: a software environment for integrated models of biomolecular interaction networks. Genome research 2003, 13:2498–2504.

82. Han H, Cho J-W, Lee S, Yun A, Kim H, Bae D, Yang S, Kim CY, Lee M, Kim E: TRRUST v2: an expanded reference database of human and mouse transcriptional regulatory interactions. Nucleic acids research 2018, 46:D380–D386.

83. Lun AT, McCarthy DJ, Marioni JC: A step-by-step workflow for low-level analysis of single-cell RNA-seq data with Bioconductor. F1000Research 2016, 5.

